# Quantitative imaging of RNA polymerase II activity in plants reveals the single-cell basis of tissue-wide transcriptional dynamics

**DOI:** 10.1101/2020.08.30.274621

**Authors:** Simon Alamos, Armando Reimer, Krishna K. Niyogi, Hernan G. Garcia

## Abstract

The responses of plants to their environment often hinge on the spatiotemporal dynamics of transcriptional regulation. While live-imaging tools have been used extensively to quantitatively capture rapid transcriptional dynamics in living animal cells, lack of implementation of these technologies in plants has limited concomitant quantitative studies. Here, we applied the PP7 and MS2 RNA-labeling technologies for the quantitative imaging of RNA polymerase II activity dynamics in single cells of living plants as they respond to experimental treatments. Using this technology, we count nascent RNA transcripts in real-time in *Nicotiana benthamiana* (tobacco) and *Arabidopsis thaliana* (Arabidopsis). Examination of heat shock reporters revealed that plant tissues respond to external signals by modulating the number of cells engaged in transcription rather than the transcription rate of active cells. This switch-like behavior, combined with cell-to-cell variability in transcription rate, results in mRNA production variability spanning three orders of magnitude. We determined that cellular heterogeneity stems mainly from the stochasticity intrinsic to individual alleles. Taken together, our results demonstrate that it is now possible to quantitatively study the dynamics of transcriptional programs in single cells of living plants.

## Introduction

Plant growth and development depends on rapid and sensitive signaling networks that monitor environmental fluctuations and transduce this information into transcriptional changes that lead to physiological adaptation. Gene regulation in plants can be extremely fast, with changes in mRNA abundance detectable in seconds to minutes, for example in response to modulations in light intensity (***Suzuki et al., 2015***; ***Crisp et al., 2017***), light quality (***Leivar et al., 2009***), the axis of gravity (***Kimbrough et al., 2004***), nutrient concentration (***Krouk et al., 2010***) or temperature (***Zandalinas et al., 2020***).

A first step toward understanding how plant transcriptional programs unfold in time and space is to quantify gene activity in individual living cells as they respond to external stimuli. Protein reporters have been used in plants to measure the dynamics of single-cell gene activity in live tissues over hours to days (***Gould et al., 2018***). However, fluorescent proteins mature at timescales that are long (>30 min) compared to the rates that characterize stress-responsive transcription (∼1 min) (***Kollist et al., 2019***), particularly in organisms grown at moderate temperatures such as plants (***Balleza et al., 2018***). In addition, protein reporter signals convolve processes such as transcription, RNA processing, RNA transport, translation, and protein degradation, often making it challenging to precisely identify where and how regulatory control is being applied along the central dogma.

In the last few years, our understanding of transcriptional regulation in animals has been transformed by techniques that quantify transcriptional activity in single cells of living embryos (***Ferraro et al., 2016; Lucas et al., 2013***; ***Gregor et al., 2014; Garcia et al., 2020***) and adult mice (***Das et al., 2018***). Here, nascent RNA is fluorescently labeled by tagging genes of interest with RNA aptamers such as MS2 or PP7 that recruit fluorescent proteins to transcriptional loci, revealing real-time transcriptional activity at the single-cell level. However, research into the equally diverse and important gene regulatory aspects of plant development and physiology has remained relatively isolated from these technological breakthroughs.

Here we bridged this technological gap by developing and implementing the PP7 and MS2 technologies for labeling nascent RNA in *Arabidopsis thaliana* (Arabidopsis) and *Nicotiana benthamiana* (tobacco). Through state-of-the-art quantitative imaging, we counted the absolute number of elongating RNA polymerase II (RNAP) molecules at individual genes and measured how this number is regulated dynamically in response to heat stress. We used this stress response in leaf tissue as a model to determine how tissue-level patterns of mRNA accumulation arise from the dynamical transcriptional behavior of individual cells. We uncovered previously unknown modes of gene regulation in plants by which tissues respond to external signals by modulating the fraction of cells engaged in transcription, but leave the single-cell transcription rate unchanged. Further, we determined how these regulatory layers give rise to a surprising level of cellular heterogeneity. The resolution afforded by PP7 and MS2 made it possible to characterize the sources of this cell-to-cell variability, revealing that stochastic processes intrinsic to individual alleles contribute to differences of three orders of magnitude in mRNA production between neighboring cells. Together, these results highlight the potential of live-imaging techniques for uncovering and quantitatively describing regulatory processes with spatiotemporal resolutions that cannot be achieved with methods such as traditional protein reporters or single-cell RNA sequencing. We envision that this approach will open new avenues of inquiry in plant cell and developmental biology.

## Results

### Establishment of the PP7 and MS2 systems for single-cell live imaging of transcription in plants

To quantitatively measure transcriptional dynamics in tobacco and Arabidopsis, we implemented an mRNA fluorescent-tagging approach previously used in animal cells in culture (***Golding et al., 2005; Chubb et al., 2006; Darzacq et al., 2007; Larson et al., 2011***), *D. melanogaster* embryos (***Garcia et al., 2013; Lucas et al., 2013***), the mouse brain (***Park et al., 2014***), and *Caenorhabditis elegans* (***Lee et al., 2019***) in which the gene of interest is tagged with tandem repeats of the PP7 DNA sequence that, when transcribed, form RNA stem-loops (Fig. 1A) (***Chao et al., 2008; Larson et al., 2011***). The PP7 loop RNA is bound by the PP7 bacteriophage coat protein (PCP) (***Chao et al., 2008***) expressed under a ubiquitous promoter. Fusing PCP to a fluorescent protein results in the fluorescent labeling of nascent RNA molecules. By virtue of the relatively slow movement of genomic loci in the nucleus and the accumulation of fluorophores in the diffraction-limited volume of the gene, sites of active transcription appear as bright fluorescent puncta over the background of nuclear PCP fluorescence in a laser-scanning confocal microscope. The fluorescence intensity of these spots reports on the number of RNAP molecules actively transcribing the gene at any given time (***Garcia et al., 2013***) and is proportional to the instantaneous rate of transcription (***Lammers et al., 2020; Bothma et al., 2014***).

**Figure 1.**
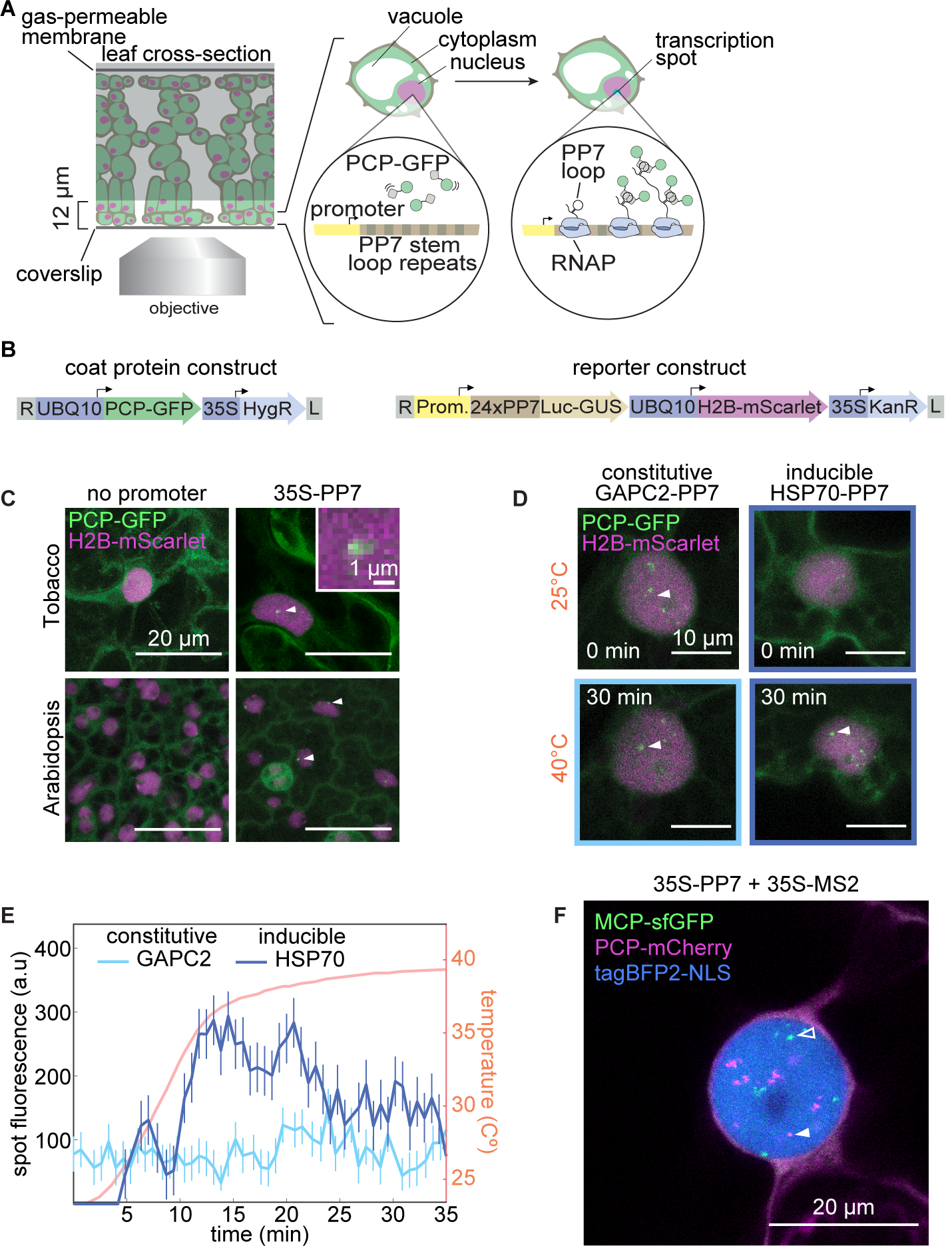
Fluorescence labeling of nascent RNA in tobacco and Arabidopsis reveals single-cell transcriptional dynamics in real time. **(A)** Schematic of the live-imaging experimental setup in leaves and diagram of the PP7 RNA labeling system. **(B)** Schematic of the constructs used in this study. (UBQ10, Arabidopsis ubiquitin 10 promoter; 35S, CaMV 35S promoter; HygR, hygromycin resistance; Luc-GUS, firefly luciferase-*/3*-glucoronidase fusion; H2B, Arabidopsis histone 2B coding sequence; KanR, kanamycin resistance; L, T-DNA left border; R, T-DNA right border). **(C)** Maximum projection of snapshots of cells expressing PCP-GFP and the reporter construct with or without the constitutive 35S promoter driving expression of the PP7-tagged Luc-GUS gene. White arrows indicate nuclear fluorescent puncta corresponding to transcription spots. Inset: magnification of PP7 fluorescence. **(D)** Maximum projection snapshots of tobacco cells expressing PCP-GFP and reporter constructs driven by the promoters of the Arabidopsis *GAPC2* and *HSP70* genes. Time under heat shock is indicated. White arrowheads indicate the fluorescent spots quantified in (F). **(E)** Fluorescence time traces of single nuclear GFP puncta in tobacco leaf epidermis cells expressing PCP-GFP and reporter constructs driven by various Arabidopsis promoters. Prior to spot detection, spots are assigned a fluorescence value of zero. Error bars represent the uncertainty in the spot fluorescence extraction (Materials and Methods). **(F)** Maximum projection snapshot of tobacco leaf epidermal cell expressing PCP-mCherry, MCP-GFP, H2B-tagBFP2, and two reporter constructs driven by the 35S promoter and tagged with PP7 (magenta) or MS2 (green). Open and closed arrowheads indicate MCP-tagged and PCP-tagged nascent RNAs, respectively.

To optimize this imaging strategy for plants, we generated two classes of constructs (Fig. 1B): (1) coat protein constructs that fuse PCP to a fluorescent protein such as GFP under a constitutive and ubiquitously expressed Arabidopsis promoter, and (2) reporter constructs that contain a neutral DNA sequence consisting of a firefly luciferase-*/3*-glucoronidase fusion with 24 PP7 stem loop repeats inserted in the 5’ end of this gene, under the control of the promoter of interest. To aid in the automated segmentation of nuclei, reporter constructs also contain a nuclear label consisting of the mScarlet red fluorescent protein (***Bindels et al., 2016***) fused to the Arabidopsis histone 2B coding region driven by a ubiquitous promoter (***Federici et al., 2012***). These two constructs confer resistance to different antibiotics, allowing sequential and combinatorial transformation into plants.

We tested this system in tobacco by simultaneously infiltrating leaves with two *Agrobacterium* strains, one strain carrying a PCP-GFP plasmid and a second strain carrying a reporter plasmid lacking a functional promoter, yielding homogeneous GFP nuclear and cytoplasmic fluorescence (Fig. 1C, top left). When the strong and constitutive 35S promoter was used to drive the reporter construct, nuclear GFP puncta became visible (Fig. 1C, top right). These results suggest that spots correspond to sites of active transcription and rule out potential PCP-GFP nuclear aggregation artifacts. Analogous results were obtained in stably transformed transgenic Arabidopsis plants (Fig. 1C, bottom).

We next sought to confirm that spot fluorescence constitutes a dynamical readout of transcriptional activity. To this end, we asked whether spot fluorescence dynamics in tobacco qualitatively recapitulate previous observations performed on the same promoters in Arabidopsis with orthogonal techniques. This comparison is made possible by the strong conservation of transcriptional regulation in plants (***Wilhelmsson et al., 2017***), in particular the heat shock response (***Mittler et al., 2012***). We measured the transcriptional activity of two well-known constitutive and heat shock-inducible Arabidopsis genes (*GAPC2* and *HSP70*, respectively (***Czechowski et al., 2005; Dong Yul Sung et al., 2001***)) before and during a heat shock treatment. GAPC2-PP7 expression was detectable at 25 °C (Fig. 1D, top left, Movie S1). The presence of multiple spots per nucleus is likely due to multiple transgene transfer events; the number of spots did not change with treatment (Figure 1D, bottom left). Further, the fluorescence of one of these spots over time did not change upon heat shock (Fig. 1E), in accordance with the constitutive expression of *GAPC2* in Arabidopsis (***Czechowski et al., 2005***). Consistent with the heat shock inducibility of the *HSP70* gene in Arabidopsis (***Dong Yul Sung et al., 2001***), HSP70-PP7 transcription was hardly detectable at 25 °C in tobacco (Fig. 1D, top right). However, upon increasing the temperature to 39 °C, multiple fluorescent puncta rapidly appeared (Fig. 1D, bottom right, Movie S1), and their fluorescence increased with time (Fig. 1E). Thus, we conclude that the PP7 system reliably recapitulates previous qualitative knowledge of transcriptional dynamics in plants.

Simultaneously tagging multiple mRNA species or multiple locations of the same mRNA species with different fluorescent proteins has revealed regulatory and physical interactions between loci and uncovered the regulation of distinct steps of the transcription cycle in cells in culture and animals (***Hocine et al., 2012; Coulon et al., 2014; Fukaya et al., 2016, 2017; Lim et al., 2018b***,a). To enable such multiplexing in plants, we also implemented the MS2 system, which is analogous and orthogonal to the PP7 system. Here, MS2 loops are specifically recognized by an MCP coat protein (MCP) (***Bertrand et al., 1998***). We tested the MS2 system in tobacco and obtained results comparable to those obtained for PP7 (Fig. S1), allowing us to track the expression dynamics of two transgenes in a single cell (Fig. 1F).

### Quantitative characterization of the PP7 system in Arabidopsis

To study transcriptional regulation at the single-cell level in populations of genetically identical leaf cells, we next generated stably transformed lines of Arabidopsis carrying PCP-GFP and a PP7 reporter construct driven by the promoter of the stress-inducible HSP101 gene. Stably transformed lines are preferable to the transient transformation in tabacco, because agroinfiltration inserts a variable number of transgenes randomly throughout the genome. A line carrying a single reporter locus (hereafter referred to as HSP101-PP7-1) was used for the following experiments unless stated otherwise; for details, see Materials and Methods: Generation of transgenic Arabidopsis lines.

A key step toward establishing PP7 as a reporter of single-cell transcriptional activity in Arabidopsis is to demonstrate that the observed spot fluorescence dynamics quantitatively recapitulate this activity. We therefore sought to cross-validate PP7 measurements with RT-qPCR quantifications of mRNA abundance in our stably transformed Arabidopsis plants. The *HSP101* mRNA is hardly detectable across vegetative tissues under standard growth conditions (***Queitsch et al., 2000***) and rapidly accumulates to high levels upon treatments inducing cytosolic protein misfolding such as heat shock (***Charng et al., 2007***). As previous experiments have shown that, upon induction, *HSP101* is expressed uniformly throughout plant tissues (***Winter et al., 2007; Jean-Baptiste et al., 2018***), we compared the average transcriptional activity of a few hundred leaf cells obtained by microscopy with that of the whole plant in bulk reported by RT-qPCR.

As expected, we did not detect actively transcribing cells in HSP101-PP7-1 plants imaged for 1 h at room temperature (Fig. S2), but shifting the microscope stage from 22 °C to 39 °C resulted in the rapid appearance of transcription spots (Fig. 2A, Movie S1). To compare the instantaneous metric of transcriptional activity reported by spot fluorescence with the number of accumulated reporter mRNA molecules captured by RT-qPCR, we converted spot fluorescence to number of produced mRNA molecules by integrating the fluorescence of all spots in the field of view over time (Fig. S3; ***Garcia et al. (2013)***.

**Figure 2.**
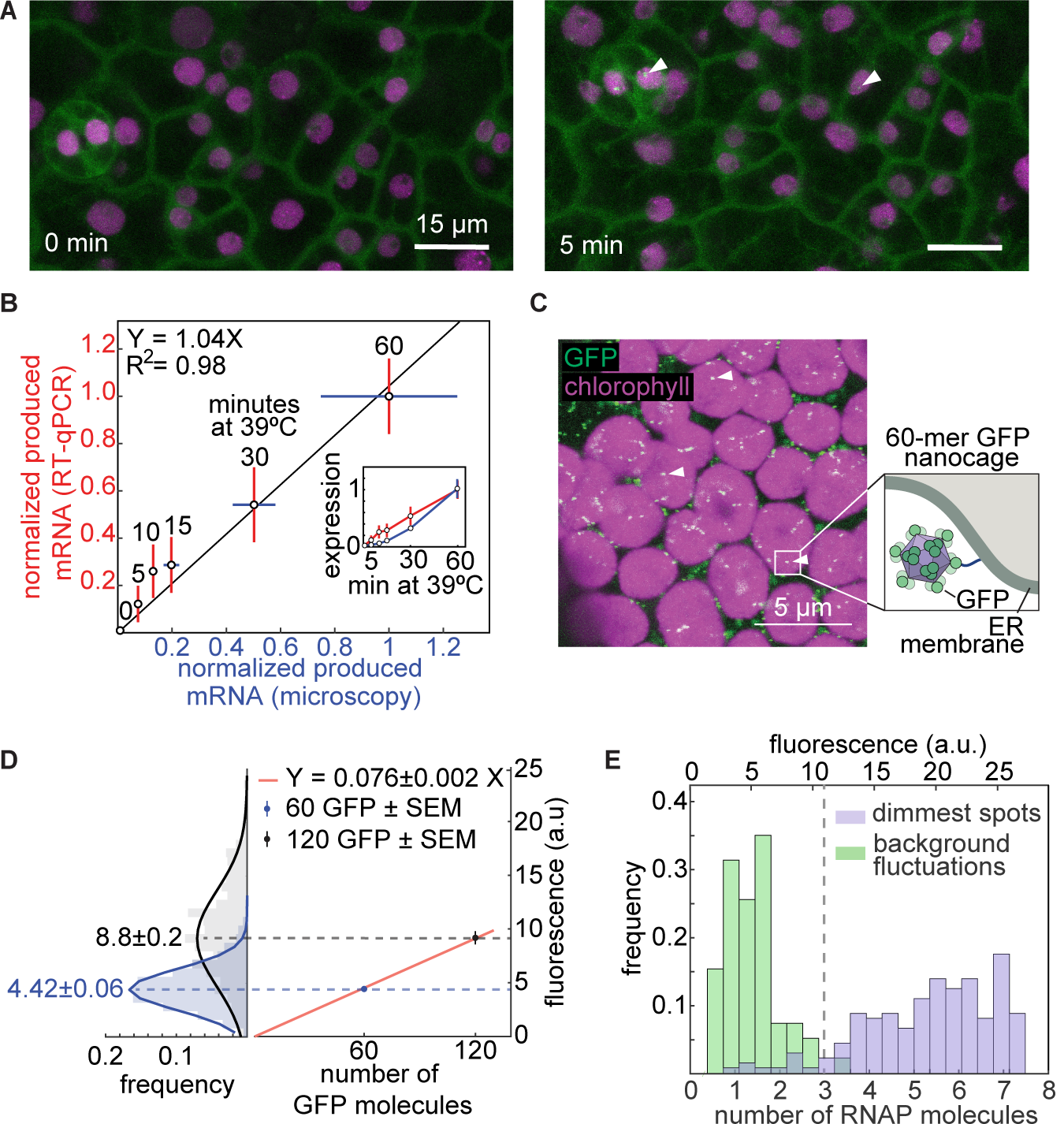
Cross validation, absolute calibration, and sensitivity of the PP7 reporter system. **(A)** Maximum fluorescence projections of leaf epidermal tissue of an Arabidopsis line stably transformed with PCP-GFP and a reporter construct driven by the HSP101 promoter under heat shock. Time stamps indicate time under heat shock. Arrowheads point to transcription spots. **(B)** Comparison between total mRNA produced as reported by RT-qPCR and PCP-GFP. PCP-GFP error corresponds to the standard error of the mean over 10 biological replicates; RT-qPCR error corresponds to the standard error of the mean (SEM) across three biological replicates. Data are normalized to each corresponding signal at 60 min. The solid black line shows a linear fit to the data going through the origin. The inset shows the normalized mean and SEM of expression level as a function of time for RT-qPCR and microscopy. **(C)** Maximum fluorescence projection of a tobacco mesophyll cell expressing a construct encoding a 60 GFP nanocage tethered to the outer membrane of the endoplasmic reticulum (ER). **(D, left)** Absolute calibration of GFP fluorescence. Histograms and Gaussian fit of single-nanocage fluorescence distributions for the 60-GFP (blue) and 120-GFP (black) nanocages transiently expressed in tobacco leaves. The mean of each distribution is shown next to each histogram. As expected, the means are related by a factor of two. **(D, right)** Mean and standard error of the mean (SEM) of the nanocage fluorescence as a function of number of GFP molecules per cage. The red line is a linear fit passing through the origin, revealing a calibration factor of 0.076 ± 0.002 a.u./GFP molecule (error reporting on the 95% confidence interval). **(E)** Histograms of the calibrated number of transcribing RNAP molecules in the dimmest three frames of the weakest half of HSP101-PP7 fluorescence time traces (purple) and their associated fluorescence background fluctuations (green). The point where the distributions overlap, at 3 RNAP molecules (vertical dashed line), can be considered the detection threshold.

Controls for GFP photobleaching ruled out the possibility that we underestimated the produced mRNA calculated by microscopy (Fig. S4). Finally, we measured HSP1010 reporter mRNA abundance by RT-qPCR using whole plants treated with heat shock (see Materials and Methods: Heat shock treatments). These measurements were strongly correlated with each other (Fig. 2B), confirming that spot fluorescence directly reports on the rate of mRNA production. This conclusion held regardless of mRNA degradation rate (Fig. S5).

While our measurements so far have shown that PP7 fluorescence is *proportional* to the number of actively transcribing RNAP molecules, it does not, by itself, report on their *absolute number*. Expressing measurements in terms of absolute number of active RNAP molecules instead of arbitrary fluorescence units is necessary for directly comparing data across microscopy setups and laboratories, and for integration with other quantitative measurements and theoretical models (***Rosenfeld et al., 2005; Cai et al., 2006; Garcia and Phillips, 2011; Garcia et al., 2013; Xu et al., 2015***). In order to turn the PP7 system into such a precision tool, we calibrated its arbitrary fluorescence units to report on the number of RNAP molecules actively transcribing the reporter gene. We followed a recently established approach to measure the fluorescence of individual GFP molecules arranged in 60-meric nanocages *in vitro* (***Hsia et al., 2016***) and *in vivo* (***Akamatsu et al., 2020***). We fused GFP to a monomer that forms these 60-meric nanocages and expressed it in tobacco leaves (Fig. 2C) to obtain a distribution of fluorescence intensity values for the resulting GFP punctae (Fig. 2D, left).

Fusing two GFP molecules to each nanocage monomer yielded the fluorescence distribution of nanocages containing 120 GFP (Fig. 2D, left). A linear fit of the means of these distributions passing through the origin shows that the mean fluorescence of 120 GFP is almost exactly twice that of 60 GFP (Fig. 2D, right), confirming the validity of this approach. The slope of this fit is an estimate of the average number of arbitrary units of fluorescence corresponding to a single GFP molecule in our microscopy setup, making it possible to report PP7 measurements in absolute units.

Our absolute calibration also provided the opportunity to determine the limits of applicability of the PP7 technology. Specifically, there is a minimum number of actively transcribing RNAP molecules below which no reliable detection is possible. Figure 2E compares histograms of the calibrated number of RNAP molecules in the weakest detectable spots and the corresponding fluctuations in background fluorescence in the data from Figure 1F. Consistent with previous measurements (***Garcia et al., 2013; Lammers et al., 2020***), these histograms overlap at approximately 3 RNAP molecules, marking the level at which PP7 fluorescent spots become undetectable. The average gene length in Arabidopsis is about 2 kbp (***The Arabidopsis Genome Iniative, 2000***) and the footprint of an elongating RNAP molecule is ≈35 bp (***Tornaletti et al., 1999***). As a result, an average gene can accommodate a maximum of 2 kbp/35 bp≈60 RNAP molecules, well above the minimum 3 RNAP molecules that constitute this detection limit. An alternative way to view this detection limit is to consider the minimum detectable rate of transcription initiation. Given an elongation rate of 1.5 kbp/min (***Ardehali and Lis, 2009***), an RNAP molecule takes 3 min to transcribe an average Arabidopsis gene. Thus, to ensure at least 3 RNAP molecules on the gene and signal detectability at any time point, transcription needs to initiate at a minimum rate of 1 RNAP/min.

### Uncovering single-cell transcriptional responses to heat shock

While static snapshots of tissues have provided profound lessons about the spatial control of transcription in animals and plants alike (***Birnbaum, 2018; Taylor-Teeples et al., 2011***), these approaches have not revealed how single-cell transcriptional dynamics dictate the temporal modulation of gene expression patterns. We sought to bridge this gap between single-cell and tissue-wide transcriptional dynamics by tracking individual nuclei and measuring the fluorescence of their corresponding transcription spot over time. To expand our range of inquiry, we generated two additional reporter lines under the control of a second heat shock-inducible promoter (HsfA2-PP7, Movie S1) or of a constitutive promoter (EF-Tu-PP7, Movie S1). In order to simplify the experiment, we imaged diploid cells of hemizygous Arabidopsis derived from the first generation of single-insertion transgenic plants (i.e., T2 individuals) such that each nucleus contained at most one spot (See Materials and Methods: Microscopy setup and image acquisition).

A striking feature of the single-cell response is the existence of a reproducible fraction of nuclei that does not show detectable expression throughout the experiment in all three assayed promoters (Fig. 3A, Fig. S6). The presence of these transcriptionally refractory cells was surprising given that endogenous *HSP101* and *HsfA2* are strongly induced and are necessary to survive heat stress in a dose-dependent manner (***Queitsch et al., 2000; Charng et al., 2007***). Similarly, as a highly expressed constitutive gene, *EF-Tu* would also be expected to be transcribed in every cell. Yet, this constitutive gene also presents a substantial fraction of refractory cells (Fig. 3A, right). Such refractory cells have also been identified in live-imaging studies of the early development of the fruit fly (***Garcia et al., 2013***; ***Lammers et al., 2020; Berrocal et al., 2020***) and in *in vitro* cultures of animal cells (***Hafner et al., 2020***).

**Figure 3.**
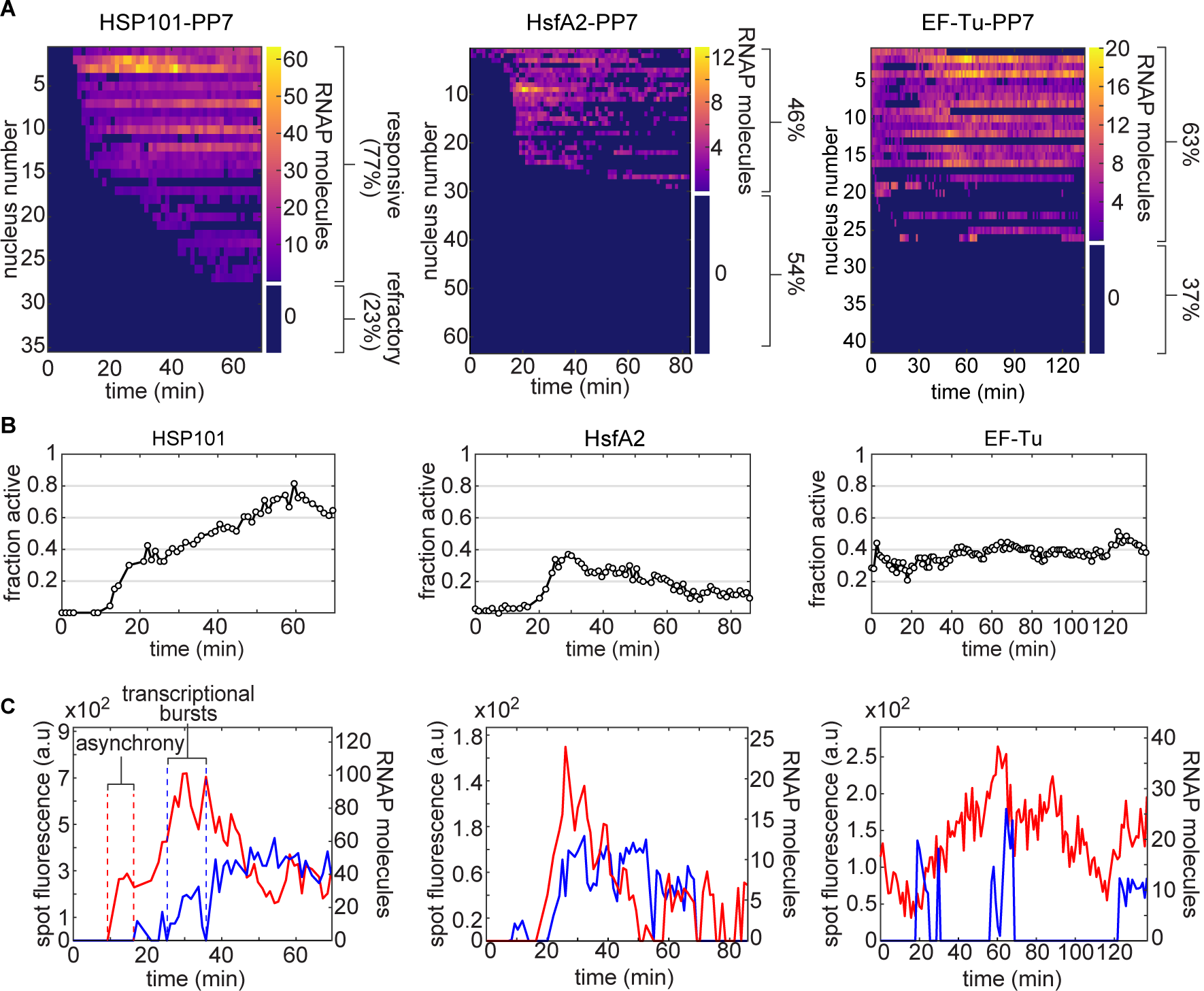
Single-cell control of transcriptional activity in response to heat shock in Arabidopsis. **(A)** Heat maps of spot fluorescence in all nuclei (rows) over time (columns) across the the field of view in HSP101-PP7-1, HsfA2-PP7-1, and EF-Tu-PP7-1 plants. Dark blue represents the absence of detectable signal. The size of the colorbar on the right of each heatmap shows the proportion of nuclei that exhibited activity in at least one frame during the experiment (>68 min) to refractory cells that presented no spots. **(B)** Instantaneous fraction of actively transcribing nuclei measured as the number of nuclei with spots divided by the total number of nuclei in the field of view. **(C)** Representative single-spot fluorescence time traces. Upon induction, transcriptional onset can occur asynchronously and transcriptional activity occurs in bursts, modulating the instantaneous fraction of transcriptionally active nuclei in (B).

To confirm that the presence of refractory cells was not an artifact of our construct or of the PP7 technology, we examined a transgenic plant containing a HSP101-GFP fusion driven by the *HSP101* promoter that fully complements the heat-susceptibility phenotype of a *hsp101* knockout (***McLoughlin et al., 2016)***. Treatment of HSP101-GFP plants with the conditions used in our PP7 experiments revealed the presence of two types of cells: cells whose fluorescence was close to that of untreated cells and highly induced cells (Figure S7). These low-fluorescence cells, which can be located right next to highly expressing ones, support the existence of transcriptionally refractory cells and the ability of the PP7 technology to detect them.

Within responsive nuclei, we also found substantial heterogeneity in the instantaneous number of actively transcribing RNAP molecules. For example, at any given time, not all responsive nuclei harbored fluorescent spots; the fraction of active nuclei is modulated in response to heat shock, but remains constant for the constitutive promoter (Fig. 3B). Interestingly, individual spots do not turn on synchronously and present periods of high transcriptional activity interspersed by periods of low to no activity (Fig. 3C). This single-cell behavior is consistent with the presence of transcriptional bursts, which have been identified across organisms and are believed to emerge from the intrinsically stochastic nature of the biochemical process of transcription (***Nicolas et al., 2017***). Interestingly, the only plant gene probed in such detail before (to our knowledge) lacked such bursts (***Ietswaart et al., 2017***).

### Tissue-wide transcriptional dynamics arise from the switch-like regulation of the instantaneous fraction of transcribing cells

How do tissue-level patterns of mRNA arise from the transcriptional activities of individual cells? Such tissue-level control could be implemented in two possible ways (***Ko, 1992***; ***Walters et al., 1995***; ***Blackwood and Kadonaga, 1998***; ***Fiering et al., 2000***). One strategy consists of modulating the single-cell rate of transcription in a graded fashion (Fig. 4A, top). Alternatively, transcriptional control could work like a switch, where the fraction of actively transcribing cells is modulated across the tissue (Fig. 4A, bottom). Several *Drosophila* enhancers invoke both strategies simultaneously (***Garcia et al., 2013***; ***Bothma et al., 2014***; ***Lammers et al., 2020***; ***Berrocal et al., 2020***). Single timepoint measurements in plants(***Turco et al., 2019***; ***Angel et al., 2011***) and live-imaging studies in cell culture (***Hafner et al., 2020***) have also provided evidence for switch-like control.

**Figure 4.**
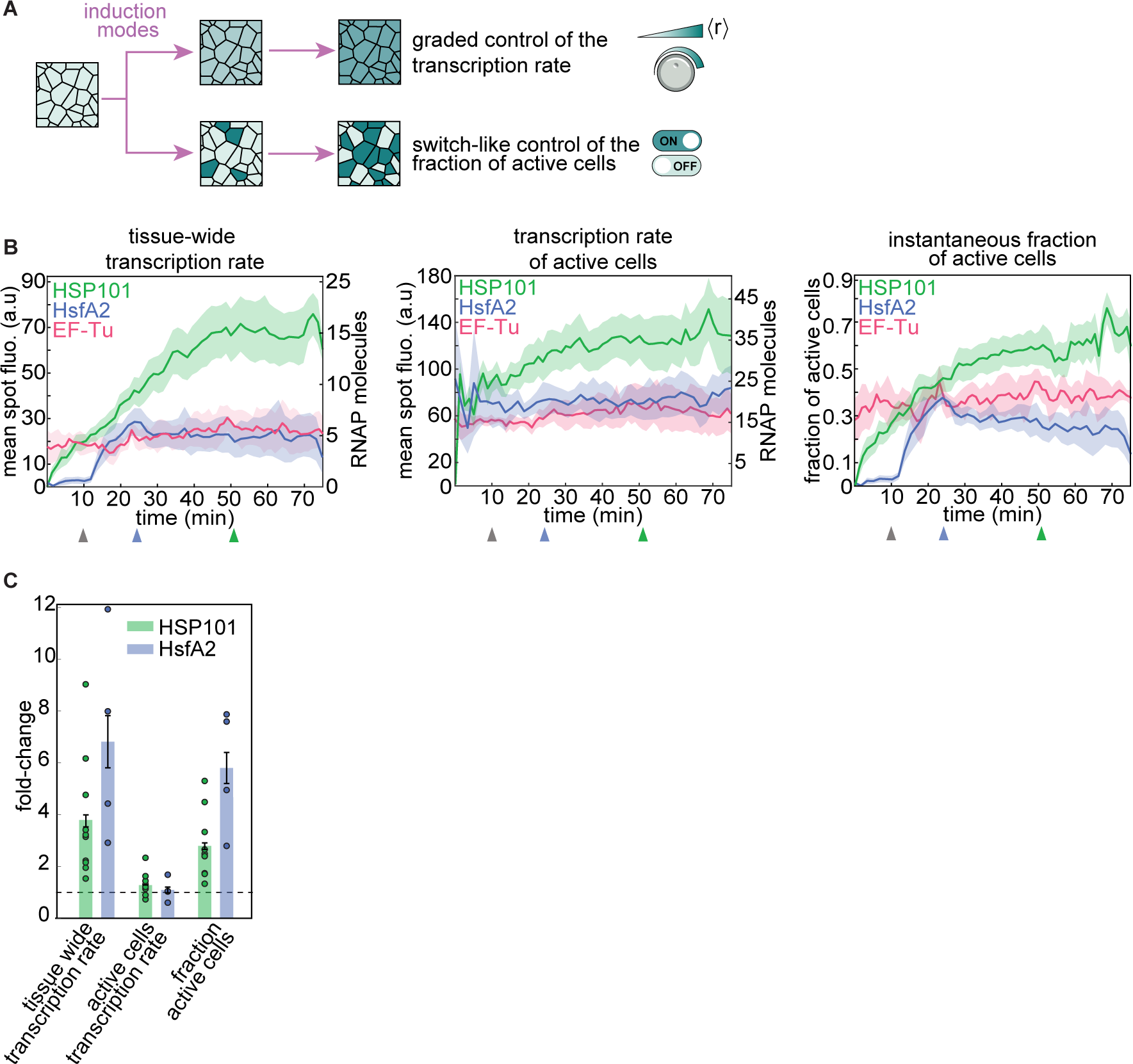
Single-cell regulatory strategies determining tissue-wide transcriptional dynamics. **(A)** Tissue-wide transcriptional control can be achieved through two non-exclusive regulatory modes: the graded modulation of the rate of transcription across cells, or the switch-like regulation of the fraction of actively transcribing cells. **(B)** Mean tissue transcription rate (left), transcription rate of active cells (middle), and instantaneous fraction of actively transcribing cells (right) for Arabidopsis lines carrying inducible promoters HSP101-PP7-1 (green) and HsfA2-PP7-1 (blue), and a line with the constitutive reporter EF-Tu-PP7-1 (red). Time *t* = 0 corresponds to the frame at which spots were first detected. **(C)** Fold-change in the mean tissue-wide transcription rate compared to the fold-change in the mean transcription rate of active cells and in the fraction of active cells, defined as the ratio between the value at its peak and at *t* = 10*min* for HSP101-PP7-1 (gray vs. green arrowheads in B) and HsfA2-PP7 (gray vs. blue arrowheads in B). The horizontal dashed line indicates a fold change of 1. (A-C, shaded regions and error bars are SEM calculated across 10, 5, and 3 experimental replicates for HSP101-PP7-1, HsfA2-PP7-1, and EF-Tu-PP7-1, respectively.)

We found that, as transcriptional induction ensues, the instantaneous fraction of cells actively transcribing increases (Fig. 3B). In addition, the level of transcription in active cells can also fluctuate (Fig. 3C). We therefore sought to determine the extent to which each regulatory strategy gives rise to tissue-wide control of the mean mRNA production rate. To this end, we expressed the total bulk transcriptional activity in terms of the quantitative contribution of each regulatory strategy as

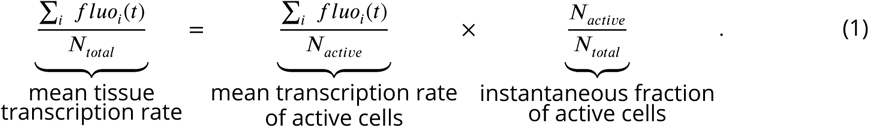

Here, *fluo*_*i*_(*t*) is the fluorescence of the *i*-th cell at time point *t, N*_*active*_ is the instantaneous number of active cells, and *N*_*total*_ is the total number of cells.

In order to determine how the resulting tissue-level transcriptional dynamics arises from the two contributions on the right side of Equation 1, we first determined the tissue-wide transcription rate at each time point by adding the fluorescence of all spots in each frame and then dividing by the total number of nuclei in the field of view (Eq. 1). The tissue-wide transcription rate of HSP101-PP7-1 and HsfA2-PP7-1 rose upon induction, while that of the constitutive EF-Tu-PP7 reporter remained constant throughout the experiment (Fig. 4B, left).

To determine whether the graded modulation of the transcription rate among active cells contributes to the mean tissue transcription rate, we calculated the mean spot fluorescence across actively transcribing cells. Further, to determine the contribution of the switch-like of regulation, we computed the instantaneous fraction of cells actively transcribing the reporter. Our calculations revealed that the temporal modulation of the transcription rate among active cells remained relatively constant throughout induction (Fig. 4B, middle). In contrast, the fraction of active nuclei was strongly modulated as a result of induction (Fig. 4B, right). Interestingly, these dynamics of the fraction of active cells were qualitatively comparable to the mean tissue transcription rate (compare Fig. 4B left and right).

To quantify the relative contribution of each of these regulatory strategies to the overall transcriptional dynamics, we measured the fold-change of each term in Equation 1. We defined this fold-change as the ratio between the value of each magnitude at peak induction (blue and green arrowheads in Fig. 4B) and at 10 min, shortly after the beginning of the response (grey arrowhead in Fig. 4B). For both heat-inducible promoters, the fold-change in the mean transcription rate across active cells was close to one (Fig. 4C). In contrast, the fold-change in the instantaneous fraction of active cells was almost identical to that of the total activity (Fig. 4C).

Thus, the duration of the treatment does not impact the rate of transcription of individual actively transcribing cells—when an individual cell transcribes, it tends to do so, on average, at a characteristic, relatively stable level regardless of induction time (Fig. S8). Instead, the time under stress modulates the tissue-wide transcription rate by increasing the probability that each individual cell engages in transcription.

### Allele-specific regulation underlies most tissue-wide heterogeneity in mRNA production in living plants

Although physiological responses occur at the tissue level, each cell must bear the phenotypic consequences of its individual gene regulatory behavior in response to stress. Studies of microorganisms and mammalian cells in culture have revealed that single-cell transcriptional responses to outside stimuli are often highly variable, leading researchers to posit that organisms possess mechanisms to buffer this “noise” or to leverage variability to drive the adoption of cellular fates that, for example, provide resistance against environmental insults such as antibiotics ***Raj and van Oudenaarden (2008)***; ***Maheshri and O’Shea (2007); Eldar and Elowitz (2010***). However, remarkably little is known about the level, functional roles, and underlying molecular mechanisms of transcriptional noise in shaping stress responses in multicellular systems like plants (***Cortijo and Locke, 2020***).

Although, on average, the rate of transcription of our heat-responsive reporters did not change with the duration of the heat treatment (Fig. 4C), at any given time point, the levels of activity across cells spanned more than two orders of magnitude (Fig. 5A). This behavior, combined with asynchronous activation (Fig. 3A) and the presence of cells that are transiently or permanently inactive transcriptionally (Fig. 4B,D), gives rise to a wide distribution in the predicted mRNA produced per cell (Figure 5B). This distribution spans more than three orders of magnitude, with a coefficient of variation (CV, standard deviation divided by the mean) of approximately 1.6.

**Figure 5.**
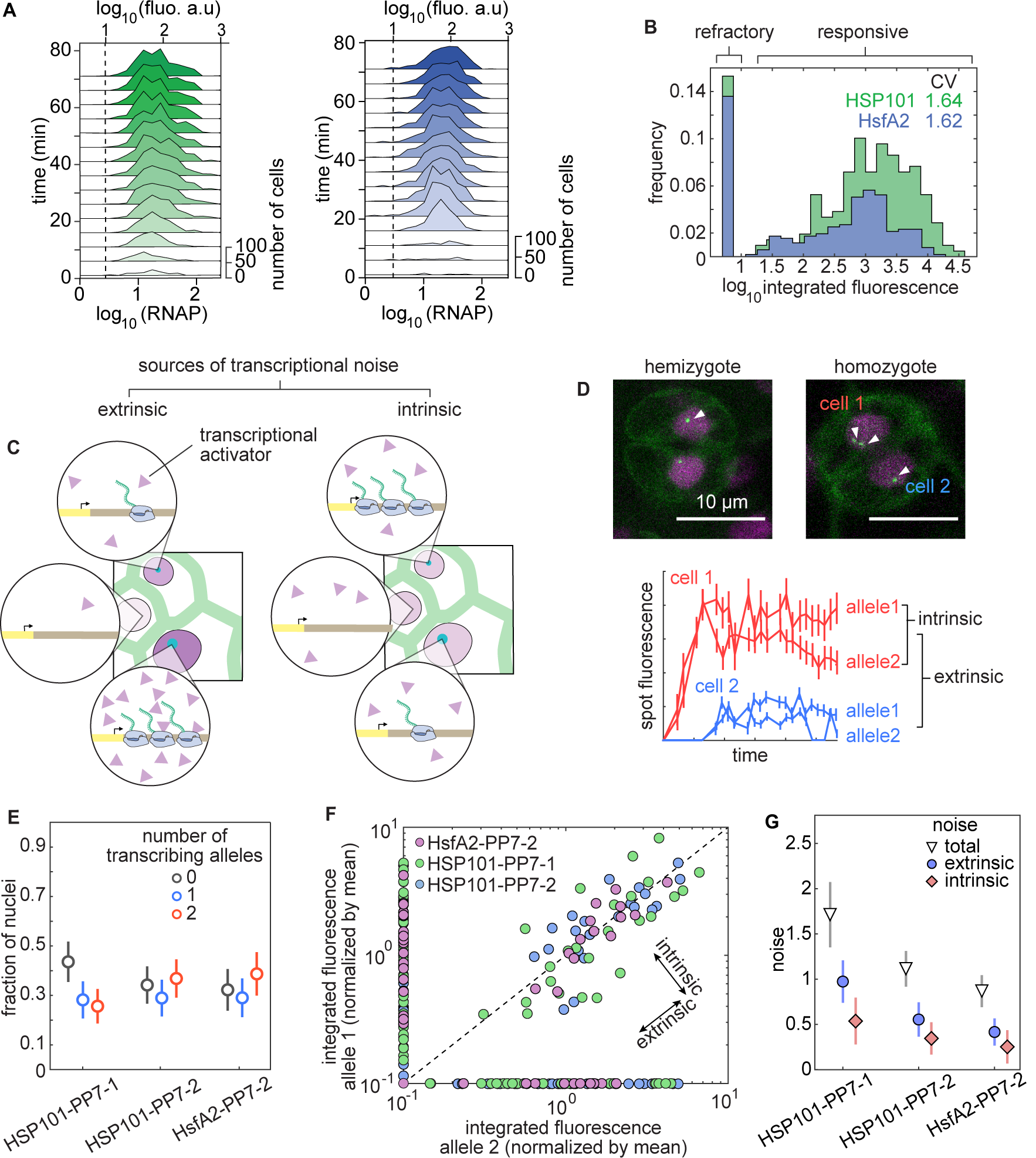
Allele-specific processes explain most of the cellular heterogeneity in produced mRNA in Arabidopsis. **(A)** Histograms of spot fluorescence over time for the combined replicates of Figure 4. The dashed line indicates the detection threshold determined in Figure2D. **(B)** Histograms of predicted total produced mRNA per cell across all replicates from Figure 4. **(C)** Schematic of extrinsic (left) and intrinsic (right) sources of transcriptional noise. Extrinsic noise arises from cellular differences in the abundance of regulatory molecules (purple triangles) while intrinsic noise captures differences among cells with identical composition. **(D)** Two-allele experiment to decompose the total transcriptional variability into intrinsic and extrinsic noise. Top: guard cells (obligate diploids) expressing HSP101-PP7. White arrowheads indicate transcription spots corresponding to one or two alleles of the reporter transgene in homologous chromosomes. In the homozygote it is possible for only one allele to be active in different cells. Bottom: spot fluorescence traces from homozygous cells shown on top. **(E)** Fraction of nuclei with zero, one, or two spots in heat shock-treated homozygous plants at the frame with the maximum number of visible spots. **(F)** Scatter plot of the integrated spot fluorescence normalized by the mean for alleles belonging to the same nucleus. Undetected spots were assigned a value of zero and plotted on the x- and y-axes. **(G)** Decomposition of the total variability in (F) into extrinsic and intrinsic components shows comparable contributions of both components to the total noise, with the intrinsic component explaining most of the variability. Error bars in (E) and (G) are bootstrapped errors.

What are the molecular sources of this cell-to-cell variability in the amount of mRNA produced (Figure 5C)? It could be the result of stochastic processes intrinsic to the allele-specific biochemical reactions that mediate transcription. In addition, a large fraction of this noise may be due to processes extrinsic to the allele itself, such as differing molecular compositions of neighboring cells. A previous measurement of gene expression noise in Arabidopsis using constitutively expressed fluorescent proteins found that extrinsic noise explains most of the cellular heterogeneity (***Araújo et al., 2017***). However, it is unclear how this noise in accumulated protein relates to transcriptional variability, and whether there are differences between constitutive and regulated promoters.

To determine whether variability is intrinsic or extrinsic to the allele, it is necessary to compare the expression of alleles belonging to the same cell with that of alleles in nearby cells (***Elowitz et al., 2002***). To make this possible, we imaged Arabidopsis individuals homozygous for the reporter, which display up to two fluorescent spots per nucleus in diploid cells (Fig. 5D, top, Movie S1). Four traces originating from two nuclei indicate that the transcriptional activity of alleles in the same nucleus can be much more similar to each other than the activity of alleles in different nuclei (Fig. 5D, bottom), suggesting a prominent role of extrinsic noise in transcriptional variability. However, our measurements also revealed that not all alleles in all nuclei are transcriptionally active: nuclei are approximately equally divided between populations presenting two, one, or even no transcription spots (Fig. 5E). The presence of nuclei with only one active allele suggests that the decision of alleles to become active is intrinsic to each allele. Thus, qualitatively, we have identified both potentially meaningful intrinsic and extrinsic contributions to the total transcriptional noise.

In order to determine the quantitative contribution of each source of variability to the single-cell distribution of mRNA produced, we followed ***Elowitz et al. (2002)*** (see Section S2 .1). To show that the results from this analysis do not depend on the number of transgene copies per insertion, we identified additional single insertion Arabidopsis lines for which we confirmed the presence of a single transgene copy per insertion locus using qPCR (see Figure S9 and associated calculations in Section S2 .2).

Figure 5F presents the integrated spot fluorescence of alleles pairs belonging to the same nucleus in homozygous plants of HSP101-PP7-1 and two additional lines with a single transgene copy per insertion. Our calculation of the noise components revealed that intrinsic sources explain most (∼ 2/3) of the variability in all of the lines tested (Fig. 5G).

In contrast to investigations of cell-to-cell variability of protein expression in constitutive promoters (***Araújo et al., 2017***), our results demonstrate that most of the cellular heterogeneity in the transcriptional response to heat shock is not due to cells having a different chemical composition. Instead, stochastic processes at the level of each individual allele explain most of the cell-to-cell differences in the amount of mRNA produced per cell. Importantly, while here we have focused on the noise in the amount of produced mRNA, further insights can be drawn from examining the sources of molecular variability in, for example, instantaneous transcriptional activity (Fig. S10).

## Discussion

Over the last few decades, it has become clear that the averaging resulting from bulk tissue sampling obscures important details about the spatial control of cellular processes in plants and animals alike. In plants, this limitation has motivated recent advances in single-cell RNA sequencing (***McFaline-Figueroa et al., 2020***). However, these measurements depend on the previous history of RNA transcription and degradation and thus obscure information about regulatory dynamics. Further, single-cell sequencing technologies tend to sacrifice spatial information. While enabling technologies to light up the process of transcription and its control in real time, in single cells or whole animals, have been developed (***Munsky et al., 2012; Tutucci et al., 2018***), plants have remained surprisingly sidelined.

Here, by implementing the PP7 and MS2 systems to fluorescently label nascent RNA molecules in plants, we have shown, to our knowledge for the first time, that it is possible to count the number of RNAP molecules actively transcribing individual alleles in single living cells of tobacco and Arabidopsis as they respond to their environment. This technical advance yielded unprecedented access to the temporal history of activity of individual alleles, making it possible to uncover distinct modes by which single-cell transcriptional activity in plants leads to tissue-wide gene expression dynamics.

Using this technique, and consistent with similar observations in other systems (***Garcia et al., 2013; Lammers et al., 2020; Hafner et al., 2020***), we discovered a fraction of transcriptionally refractory cells that do not transcribe regardless of induction conditions (Fig. 4D). Single-molecule RNA FISH experiments in Arabidopsis roots found that at any given time ≈20% of cells are transcriptionally inactive for the constitutively expressed *PP2C* gene (***Duncan et al., 2016***). However, unlike the live-imaging approach developed here, single-molecule RNA FISH relies on fixed samples; it cannot determine whether this inactive state was transient or stable.

We also found that tissue-wide transcriptional induction dynamics are the result of the temporal modulation in the fraction of cells that switch to a transcriptionally active state, and not of the graded control of the transcription rate of active cells (Fig. 4C). This form of regulation has been hypothesized to be at play in the regulation of the FLC gene in response to temperature (***Angel et al., 2011***) and in the commitment to xylem cell fate in response to the VND7 transcription factor (***Turco et al., 2019***). Using our technologies, it should now be possible to directly test these models.

Gene expression can vary significantly from cell to cell in microbial and animal species (***Raj and van Oudenaarden, 2008***). By making it possible to measure cell-to-cell transcriptional variability in real time in living plant cells, we confirmed that plants are no exception to this widespread presence of transcriptional variability. The single-locus resolution of our method allowed us to determine that cell-to-cell variability in mRNA production arises mainly from stochastic processes instrinsic to each allele (Fig. 4G). Studies in *in-vitro* cell cultures have found that gene-expression noise can have profound consequences for cellular survival (***El Meouche et al., 2016;* Shaffer et al., 2017**); however, the role of transcriptional noise in plant stress responses remains an open question (***Cortijo and Locke, 2020; Roeder, 2018***). We envision that the strategy applied here to systematically dissect transcriptional heterogeneity in Arabidopsis and tobacco will shed light on this interplay between transcriptional variability and stress response. Further, it will be interesting to examine how some unusual aspects of plant cell biology and genetics can buffer transcriptional noise. For example, cytoplasmic connections could play a role in short-range sharing of gene products (***Faulkner, 2018***), averaging out extrinsic noise; multiple genome copies per nucleus in mature plant cells may provide further opportunities to average out intrinsic noise across alleles (***Lee et al., 2019***). Similarly, we speculate that the conspicuous retention of large numbers of seemingly redundant gene paralogs in plants may also help buffer intrinsic fluctuations in individual genes (***Li et al., 2015***).

Our approach requires access to a confocal microscope and to transgenesis tools, and should therefore be relatively easy to apply to many biological problems in plant development and physiology. However, imaging deep into tissues with the resolution necessary to resolve diffraction-limited spots remains a challenge, particularly in plants. Advances such as multiphoton imaging, lattice light-sheet microscopy, and adaptive optics will overcome this limitation (***Liu et al., 2018***).

Lacking single-polymerase resolution currently limits the applicability of MS2 and PP7 to genes transcribed at relatively high rates. A transcription initiation rate of 1 RNAP/min, corresponding to our detection limit of 3 elongating RNAP molecules on an average Arabidopsis gene, could be sufficient to sustain slow transcriptional processes operating at long developmental timescales. For example, the FLC gene, a key seasonal developmental regulator in Arabidopsis is rarely occupied by more than one elongating RNAP at a time (***Ietswaart et al., 2017***) which may explain why previous attempts at visualizing nascent FLC mRNAs in live Arabidopsis plants have failed (***Wu et al., 2016***). A growing interest in live imaging of transcription combined with advances in fluorophore chemistry (***Iwatate et al., 2020***) as well as in the PP7 and MS2 technologies themselves (***Wu et al., 2012***) offer hope for breaking this detection threshold.

It will undoubtedly be of interest to correlate the activities of genes by visualizing their transcription simultaneously. This multiplexing is already possible for two genes using MS2 and PP7. A third color could be added by implementing interlaced MS2 and PP7 loops (***Hocine et al., 2012***). To further extend the palette, it should be possible to engineer other orthogonal RNA-binding proteins-RNA aptamer pairs (***Daigle and Ellenberg, 2007; Katz et al., 2018***).

Finally, and more generally, the random integration of transgenes in plants makes it challenging to dissect the role(s) of regulatory sequences at their endogenous genomic locations. Delivery of DNA with CRISPR/Cas9 or site-specific recombinases promises to unleash the potential of quantitative reporters of gene expression.

In this study, we focused on a simple step in the plant’s use of temperature as a signaling input. More complex treatments have been previously used to show that plants can mount specific responses to inputs, such as memory in response to pulses of heat shock (***Charng et al., 2007***) and nonlinear integration of combinations of high light and temperature stress (***Zandalinas et al., 2020***). By administering experimental treatments while simultaneously measuring their effects on gene regulation, it will be possible to determine how these operations are performed at the cellular level. In addition, the sub-nuclear resolution of nascent RNA tagging could make it possible to resolve long-standing issues in plant signaling, such as the role of protein aggregates or “nuclear speckles” that are pervasive in light-responsive signaling pathways in plants (***Ronald and Davis, 2019***).

In conclusion, by enabling the measurement of transcription at high spatiotemporal resolution, the PP7 and MS2 methods introduced here close a critical technological gap in plant biology. These new technologies open new avenues of inquiry and will make it possible to quantitatively interrogate transcriptional control in living plants and to engage in the discourse between theory and experiment that has characterized the study of gene regulation in single cells and animal tissues over the last two decades.

## Materials and Methods

### Plasmids and Agrobacterium strains

All plasmid sequences used in this study can be accessed from a public Benchling folder. Plasmids will be made available at Addgene. All vectors were based on pCambia derivatives (***Hajdukiewicz et al., 1994***) and transformed into the GV3101::pMP90 Agrobacterium strain by electroporation. Plasmids confering Kanamycin resistance in plants (i.e reporter constructs) were based on pCambia2300. Plasmids confering Hygromycin resistance in plants (i.e PCP, MCP and nanocages constructs) were based on pCambia1300. A list of the plasmids used in this study can be found in table S1. The Arabidopsis gene identifiers associated with genomic sequences used in these plasmids are listed in table S3.

### Plant growth conditions

*Nicotiana benthamiana* (tobacco) plants were grown in a greenhouse under natural light conditions prior to agroinfiltration. Following infiltration, tobacco plants were kept under 30 *µ*E of constant light. Arabidopsis plants used for experiments were grown in 1/2 strength MS agar containing 50 *µ*g/*µ*l of Kanamycin under short day conditions (8 hours of 30 *µ*E light per day) for four to six weeks prior to imaging.

### Agroinfiltration

Agrobacterium glycerol stocks were streaked on LB plates containing 50 *µ*g/*µ*l Kanamycin and 50 *µ*g/*µ*l Gentamycin. Fresh colonies were grown overnight in liquid LB containing the same antibiotic concentrations, spun down and resuspended in an equal volume of infiltration buffer (10 mM MES pH5.6, 10 mM MgCl_2_, 150 *µ*M Acetosyringone). Cells were incubated for 2-4 hours in infiltration buffer shaking at room temperature after which the cultures were diluted 1:3 to an OD_600_ of approximately 0.3. In experiments that required combining strains, coat protein and reporter strains were mixed in a 3:1 ratio (the exact ratio does not qualitatively affect the results). In PP7 and MS2 experiments, infiltrated leaves were imaged approximately 2 days after infiltration. For absolute calibration experiments, plants were imaged 12-18 hours after infiltration.

### Generation of transgenic Arabidopsis lines

To generate lines carrying both PCP-GFP and PP7 reporters we followed a sequential transformation approach. We first selected PCP-GFP lines in 35 *µ*g/ml of Hygromycin and kept lines exhibiting moderate levels of fluorescence and no obvious growth phenotype. Next, we transformed T1 or T2 PCP-GFP individuals with PP7 reporter Agrobacterium strains and selected transformants in 50 *µ*g/ml Kanamycin and 35 *µ*g/ml Hygromycin. Individuals T1 for the PP7 construct were screened for nuclear mScarlet fluorescence and presence of transcription spots matching previous knowledge about the activity of the corresponding endogenous gene. In all cases, to select for antibiotic resistance we followed the protocol by ***Harrison et al***. (***2006***). A list of the lines used in this study can be found in table S2.

### Determining the number of unlinked reporter transgene insertions

To select lines carrying a single insertion reporter locus we plated approximately 60 T2 seeds in MS plates containing Kanamycin and counted the ratio of survivors. This ratio was divided by the survival ratio in plates containing no antibiotics. A *χ*^2^ test was used to determine whether the product of these two ratios was statistically different from the expected ratio of 3/4. To confirm the absence of two or more unlinked reporter loci we examined transcription spots in guard cells. Unlike other leaf cell types, these cells are exclusively diploid (***Melaragno et al., 1993***) and therefore the presence of a single spot per guard cell nucleus in a T1 individual confirms the absence of unlinked insertions.

### Heat shock treatments

To control the sample temperature in the microscope stage we used an OkoLabs H101-LG temperature chamber calibrated to achieve a maximum of ≈39°C. The temperature experienced by the sample was determined once using an electronic probe. The heat shock treatment used for the RT-qPCR experiment in Figure 2A was performed as follows: whole 4-6 week-old plants were placed in 1.7 ml plastic tubes containing 200 *µ*l of water. The sample corresponding to time = 0 minutes was immediately taken out of the tube, quickly tapped dry, transferred to a new tube containing silica beads and frozen in liquid nitrogen. The rest of the samples were transferred to a 39°C heat block and removed at set times. Plants were then quickly tapped dry and frozen in liquid nitrogen.

### Microscopy setup and image acquisition

In tobacco experiments, a piece of infiltrated leaf spot was mounted in water between a glass slide and a glass coverslip with the abaxial (bottom) side facing the objective. To image the GFP nanocages in mesophyll cells, the abaxial epidermis was first removed. In Arabidopsis experiments, full 2-4 day old leaves from 4-6 week old plants were mounted in tap water between a gas permeable cellophane membrane (Lumox film; Starstedt) and and a glass coverslip with the adaxial (top) side facing the objective. All samples were imaged close to the base of the leaf blade immediately after mounting. All data was taken in a Leica SP8 confocal microscope with a white light laser using a 63X oil objective. The dimensions of the field of view were 92.26 x 46.09 *µ*m using 1052 x 512 pixels, resulting in a pixel size of 90 nm. Z stacks consisting of 25 slices of 0.5 *µ*m each were taken every 60 seconds accumulating fluorescence 3 times over lines. The beginning of each stack was set to the upper-most nucleus in the leaf epidermis. For GFP, excitation 488 nm and emission 498-559 nm. For mScarlet, excitation 569 nm, emission 579-630 nm. For Chlorophyll, excitation 488 nm, emission 665-675 nm. To ensure quantitative consistency across experiments, the 488 nm laser power at was calibrated to 10.5 *µ*W (≈5% laser power) at the beginning of each imaging session using a power meter. The percentage intensity of the 569 nm laser line was kept consistent across experiments at 5%.

### RT-qPCR

Total RNA was extracted using the Quiagen RNeasy kit following the manufacturer instructions. Reverse transcription was performed using the Qiagen Omniscript kit with a primer mix of random 10mers (10 *µ*M final concentration) and 15mer oligo dT primers (1 *µ*M final concentration). mRNA abundance was calculated by the delta CT method. Primers for endogenous HSP101 were 5’GGTCGATGGATGCAGCTAAT and 5’CTTCAAGCGTTGTAGCACCA from ***Yoshida et al***. (***2011***). Primers for the Actin2 standard were 5’CGCTCTTTCTTTCCAAGCTCAT and 5’GCAAATCCAGCCTTCACCAT from ***Liu and Ma (2011***). Primers for the reporter mRNA were 5’GGGTTCATCAGAGTGCCAGAG and 5’AGGCAGAGCGACACCTTTAG. A negative control was performed under identical conditions replacing the RT enzyme with water.

### Image analysis: spot luorescence and tracking

Raw image stacks of the coat protein channel were used to identify fluorescent punctae corresponding to transcription spots using the ImageJ implementation of the 3D Trainable Weka Segmentation toolbox (***Arganda-Carreras et al., 2017***). Following ***Lammers et al***. (***2020***), after segmentation, spots in each z-slice were fitted to a 2D Gaussian. The z-slice with the largest Gaussian amplitude was selected for the spot fluorescence calculation. Spot fluorescence corresponds to the sum of pixel intensity values in a circle with a radius of 1.08 *µm* centered around the center of the fitted Gaussian minus the background fluorescence offset. Per pixel background fluorescence is calculated from the baseline of the spot Gaussian fit. The imaging error associated with each spot trace is determined from the fluctuations over time in the per pixel offset multiplied by the spot integration area. A spline is fitted to the offset time trace to calculate offset error as the standard deviation around this spline. False negative and false positive spots were corrected manually.

### Image analysis: nuclear segmentation and spot tracking

Maximum intensity projections of the nuclear marker channel were used for nuclear segmentation using the ImageJ implementation of the 2D Trainable Weka Segmentation toolbox (***Arganda-Carreras et al., 2017***) or a custom-written Matlab pipeline. False negative and false positive nuclei were then manually corrected. Spots were assigned to nuclei based on physical overlap. Tracking of spots over time was based on nuclear tracking and manually corrected whenever errors were found.

### Image analysis: nucleus luorescence

A binary mask of segmented nuclei was applied to the PCP-GFP or Histone 2B-mScarlet channel. For each z-slice and frame, the mean fluorescence across pixels within each nucleus area was calculated. Then, for each frame, we took the intensity of the z-slice with the maximum mean fluorescence as the metric for fluorescent protein concentration in that given frame.

### Determining transgene copy number by qPCR

Genomic DNA was extracted from leaf tissue using CTAB and phenol:chlorophorm precipitation. Primers used to amplify the reporter transgene were 5’gacgcaagaaaaatcagagagatcc and 5’ggtttc-tacaggacggaccatacac. Primers used to amplify a region near the *Lhcb3* gene used as an internal genomic control were 5’acaggtttggtcaagtcaattacga and 5’atggtttccatgaatactgaacacg. The final concentration of genomic DNA per reaction was 0.75 ng. For a more detailed explanation of the calculations and controls related to this experiment see Section S2 .2.

### Absolute calibration using nanocages

Tobacco leaves were infiltrated with agrobacterium strains containing plasmids where the promoter of the Arabidopsis UBC1 gene (1138bp upstream of the AT1G14400 start codon) was used to drive the 60mer monomer fused to either one or two mGFP5 coding sequences. The N terminus of the rabbit Cytochrome P450 CII1 was added as an N terminal tag to target the protein fusions to the cytosolic side of the ER. Mesophyll cells were imaged no later than 15 hours after infiltration since longer incubation resulted in the appearance of large GFP aggregates.

## Acknowledgments

We would like to thank Rob Phillips, Setsuko Wakao, Christopher Gee and Avi Flamholz for comments on the manuscript, Allison Schwartz, Jose O’brien and Fernan Federici for sharing plasmids, and Albert Lin and Jonathan Liu for their feedback regarding calculations. HGG was supported by the Burroughs Wellcome Fund Career Award at the Scientific Interface, the Sloan Research Foundation, the Human Frontiers Science Program, the Searle Scholars Program, the Shurl and Kay Curci Foundation, the Hellman Foundation, the NIH Director’s New Innovator Award (DP2 OD024541-01), and an NSF CAREER Award (1652236). KKN is an investigator of the Howard Hughes Medical Institute.

## Supplementary Information

### S1 Biological material

**Table S1.**

| Plasmids |  |  |  |
| --- | --- | --- | --- |
| Plasmid Name | Codes for | Function | Addgene |
| UPG | AtUBQ10p::PCP-mGFP5<br>(hyg resistance in plants) | Ubiquitous expression of<br>PCP-GFP fusion |  |
| UPmCh | AtUBQ10p::PCP-mCherry<br>(hyg resistance in plants) | Ubiquitous expression of<br>PCP-mCherry fusion |  |
| UMsfG | AtUBQ10p::MCP-sfGFP<br>(hyg resistance in plants) | Ubiquitous expression of<br>MCP-sfGFP fusion |  |
| AL13Rb | PP7-Gus-Luc +<br>AtUBQ10p::H2B-mScarlet<br>(kan resistance in plants) | Promoterless PP7 reporter<br>and red nuclear marker |  |
| AL12R | AtUBQ10p::H2B-mScarlet<br>+ PP7-Gus-Luc (kan resis-<br>tance in plants) | Promoterless PP7 reporter<br>and Histone-mScarlet RFP<br>nuclear marker |  |
| AL13Rb-35S | 35S-PP7 reporter in AL13Rb | Reports on 35S promoter<br>activity and labels nuclei |  |
| AL13Rb-GAPC2 | GAPC2-PP7 reporter in<br>AL13Rb | Reports on Arabidopsis<br>GAPC2 promoter activity<br>and labels nuclei |  |
| AL12R-HSP70 | HSP70-PP7 reporter in<br>AL12R | Reports on Arabidopsis<br>HSP70 promoter activity<br>and labels nuclei |  |
| AL13Rb-HsfA2 | HsfA2-PP7 reporter in<br>AL13Rb | Reports on Arabidopsis<br>HsfA2 promoter activity<br>and labels nuclei |  |
| AL12R-EF-Tu | EF-Tu-PP7 reporter in<br>AL12R | Reports on Arabidopsis EF-Tu promoter activity and la-<br>bels nuclei |  |
| AL12R-HSP101 | HSP101-PP7 reporter in<br>AL12R | Reports on Arabidopsis<br>HSP101 promoter activity<br>and labels nuclei |  |
| UtB2N7 | AtUBQ10p::tagBFP2-NLS | nuclear localized blue fluo-<br>rescent protein marker |  |
| UBC1cer60G | AtUBC1::60mer-mGFP5 | Weak ubiquitous expres-<br>sion of an ER-targeted<br>60mer monomer fused to<br>mGFP5 |  |
| UBC1cer120G | AtUBC1::mGFP5-60mer-mGFP5 | Weak ubiquitous expres-<br>sion of an ER-targeted<br>60mer monomer fused to<br>two mGFP5 |  |

**Table S2.**

| Arabidopsis lines generated in this study |  |  |
| --- | --- | --- |
| Name | Transgenes (refer to the 'Plasmids' table) | Usage |
| UPG-6 | UPG | For transformation with reporter constructs |
| UPG-9 | UPG | For transformation with reporter constructs |
| AL13Rb-35S | UPG and AL13Rb-35S | Image 35S promoter activity in Figure 1 |
| AL12R-HSP101-1 | UPG and AL12R-HSP101 | Image AtHSP101 promoter activity in Figures 2 to 5 |
| AL13Rb-HSP101-2 | UPG and AL13Rb-HSP101 | Image AtHSP101 promoter activity in Figure 5 |
| AL13Rb-HsfA2-1 | UPG and AL13Rb-HsfA2 | Image AtHsfA2 promoter activity in Figures 3 and 4 |
| AL13Rb-HsfA2-2 | UPG and AL13Rb-HsfA2 | Image AtHsfA2 promoter activity in Figure 5 |
| AL12R-EF-Tu-1 | UPG and AL12R-EF-Tu | Image AtEF-Tu promoter activity in Figures 3, 4 and Fig. S4 |

**Table S3.**

| Arabidopsis Gene Identifiers |  |  |
| --- | --- | --- |
| Gene abbreviation | Gene name | AGI |
| UBQ10 | Polyubiquitin 10 | AT4G05320.2 |
| H2B | Histone 2B | AT5G22880.1 |
| GAPC2 | Glyceraldehyde-3-phosphate dehydrogenase C2 | AT1G13440.1 |
| HSP70 | Heat shock protein 70 | AT3G12580.1 |
| UBC1 | Ubiquitin carrier protein 1 | AT1G14400.1 |
| HSP101 | Heat shock protein 101 | AT1G74310.1 |
| HsfA2 | Heat shock transcription factor A2 | AT2G26150.1 |
| EF-Tu | GTP binding Elongation factor Tu family protein | AT1G07920.1 |

### S2 Calculations

#### S2 .1 Decomposition of total variability into extrinsic and intrinsic noise

In this section we derive the formulas for the total, intrinsic and extrinsic noise (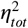, 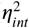, and 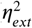, respectively) based on the two-reporter approach developed by ***Elowitz et al. (2002)***. As noted by ***Hilfinger and Paulsson (2011)*** and explained at length by ***Fu and Pachter (2016)***, these expressions stem from the law of total variance, which states that, for a random output variable *A* and a random input variable *X*, the total variance of *A* can be decomposed as the sum

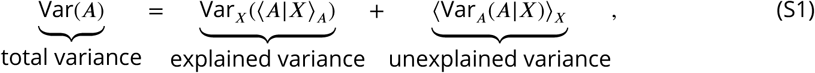

where the subscripts *X* or *A* indicate that the average or the variance is taken over different values of *X* or *A*, respectively.

Applied to the problem of gene expression variability, *A* represents the expression level of the gene of interest and *X* corresponds to the cellular state indicating, for example, the concentration in each given cell of all molecules that affect the expression of that gene such as RNAP. The first term on the right-hand side of Equation S1 is referred to as the *explained variance* and captures how much the average value of *A* varies across different values of *X*. The second term is referred to as the *unexplained variance* and captures how much the expression of *A* varies in cells that share the same value of *X*.

Because the identity and values of *X* are typically not known and/or not experimentally accessible, ***Elowitz et al. (2002)*** devised a two-reporter system to determine the explained and unexplained components of the total normalized variance, which they termed extrinsic 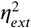 and intrinsic 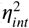 noise, respectively. In this approach, each cell has two identical but distinguishable alleles of the gene of interest. In their statistical model, these two alleles are identical in all respects meaning that their distribution over cells and over time are the same. For the purpose of this derivation, let us call *A*_*i*_ and *B*_*i*_ the expression level of each allele in the *i*-th cell and normalize *A* and *B* to their means such that

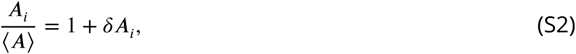

where *δA*_*i*_ is the fractional deviation of the expression level *A*_*i*_ from the mean ⟨*A*⟩. Similarly, for B we normalize to

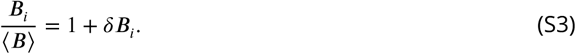

In the following calculations we will make use of the measurable quantities *δ A*_*i*_ and *δ B*_*i*_ to eliminate the unknown quantity *X* from Equation S1. We start by deriving an expression for 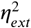 here as the explained component of the total variance of the normalized *δ A* distribution

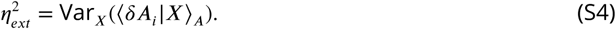

Note that, since *X* is a random variable, so is (*δ A*_*i*_|*X*)_*A*_, and we can write its variance as defined

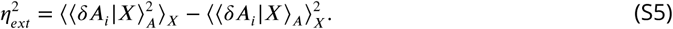

Because both alleles are identical, ⟨ *δ A*_*i*_ | *X* ⟩_*A*_ is equal to ⟨ *δ B*_*i*_ | *X* ⟩_*B*_, which allows us to write Equation S5 as

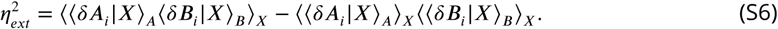

Note that, in this model, the variability in the values of *A*_*i*_ and *B*_*i*_ for cells with the same *X* are independent of each other since we assume that they are not explained by *X*. Because of this independence, ⟨*A*_*i*_⟩ ⟨*B*_*i*_⟩= ⟨*A*_*i*_*B*_*i*_⟩ for a given *X*. Applied to the first term in Equation S6, the extrinsic noise can be written as

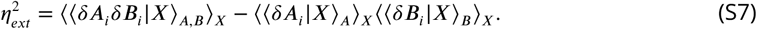

We now note that the double angle brackets in the first term in the righthand side of Equation S7 call for averaging the value of *δA*_*i*_*δB*_*i*_ in cells with the same *X* and then averaging again over all possible values of *X*. Similarly, the second term in the equation calls for averaging over *A*_*i*_ or *B*_*i*_ for a given *X*, and then averaging over *X*. This allows us to eliminate *X* in the equation and simplify our expression to

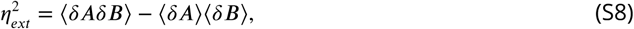

which is the definition of covariance. Thus,

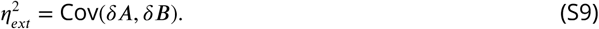

This makes intuitive sense, as the model assumes that, since *A* and *B* are identical genes that respond to *X* in the exact same way, the variance in the expression of *A* that is explained by *X* is identical to the variance in the expression of *B* that is explained by *X*. As a result, the extrinsic noise measures how *A* and *B* coordinately vary across cells.

We now turn our attention to the derivation of the intrinsic noise, which we define as the unexplained component of the variance in the normalized *A* distribution, namely

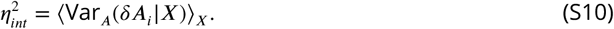

Replacing the unexplained variance in Equation S1 with 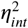, the explained variance by its formulation as extrinsic noise from Equation S9, and rearranging leads to

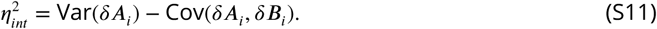

Because this equation does not involve *X* we don’t need the subscripts anymore: all variances are calculated across values of *δA* and *δB*. We now note that the total variance of *δA* and *δB* must be the same since they have the same distribution over cells and over time. Therefore we are allowed to express the first term in the right-hand side of Equation S11 as the average variance of the *δ A*_*i*_ and *δ B*_*i*_ distributions

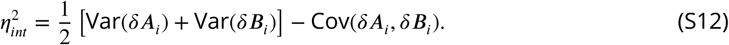

Rearranging Equation S12 leads to

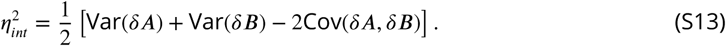

Now, using the identity stating that the variance of a sum is the sum of the variances minus their covariance, Equation S13 becomes

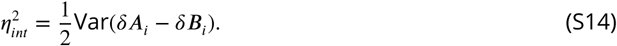

Finally, we define the total noise 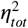 as the total variance of the normalized *δA*_*i*_ distribution. As noted before, because the distributions of *δA*_*i*_ and *δB*_*i*_ are identical, so are their variances. Therefore, the total noise can be calculated from the average

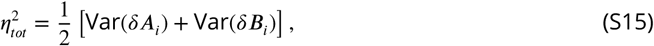

which satisfies

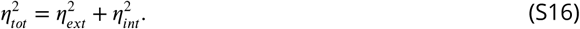

Note that, here, we considered *δ A* loosely as the “expression level” of gene *A*. This analysis can be applied to any metric of gene expression such as the instantaneous transcription rate, or the total amount of produced mRNA.

#### S2 .2 Determining transgene copy number by qPCR

In this section, we present our calculation for determining the number of transgene insertions from the Δ*Cr* values resulting from qPCR taking the amplification efficiency into account. Given a starting number of DNA molecules *N*_0_, the total number of molecules after *C* amplification cycles is given by

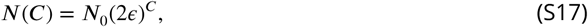

where *ϵ* corresponds to the amplification efficiency, or the fraction of molecules that are duplicated in each cycle. The number of amplification cycles *CT* that takes to amplify the number of DNA molecules from *N*_0_ to *N*_*ct*_ can be described by

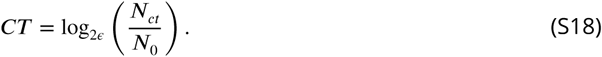

Changing the logarithm base and rearranging leads to

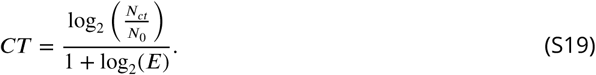

We now define an amplification efficiency constant *K* as

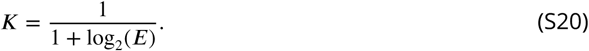

Equation S19 then becomes

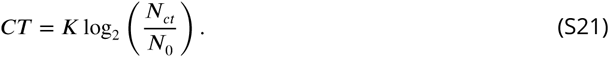

To experimentally obtain *K* (and therefore *ϵ*), we perform qPCR on serial dilutions of template DNA, thus varying *N*_0_. We then plot *CT* as a function of the log_2_ of the template concentration in order to obtain *K* from the slope (Fig. S9A,B). We used genomic DNA from a transgenic Arabidopsis plant to perform this amplification on the PP7 transgene as well as on an internal control genomic sequence. We measured both PCR reactions to have an efficiency of *K* = 1 within experimental error. As a result, we can determine the ratio between the initial number of transgene molecules 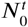 and the initial number of internal control molecules 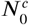 by calculating the Δ*Cr*

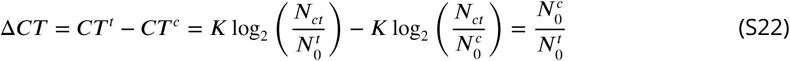

If the transgene occurs in a single insertion locus containing a single transgene copy per insertion, then in a T1 individual

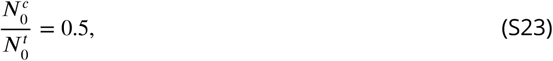

which corresponds to a Δ*Cr* value of −1. Using this approach we were able to identify transgenic Arabidopsis individuals with a single insertion locus containing a single transgene insertion (Fig. S9C).

### S3 Supplementary Figures

**Figure S1.**
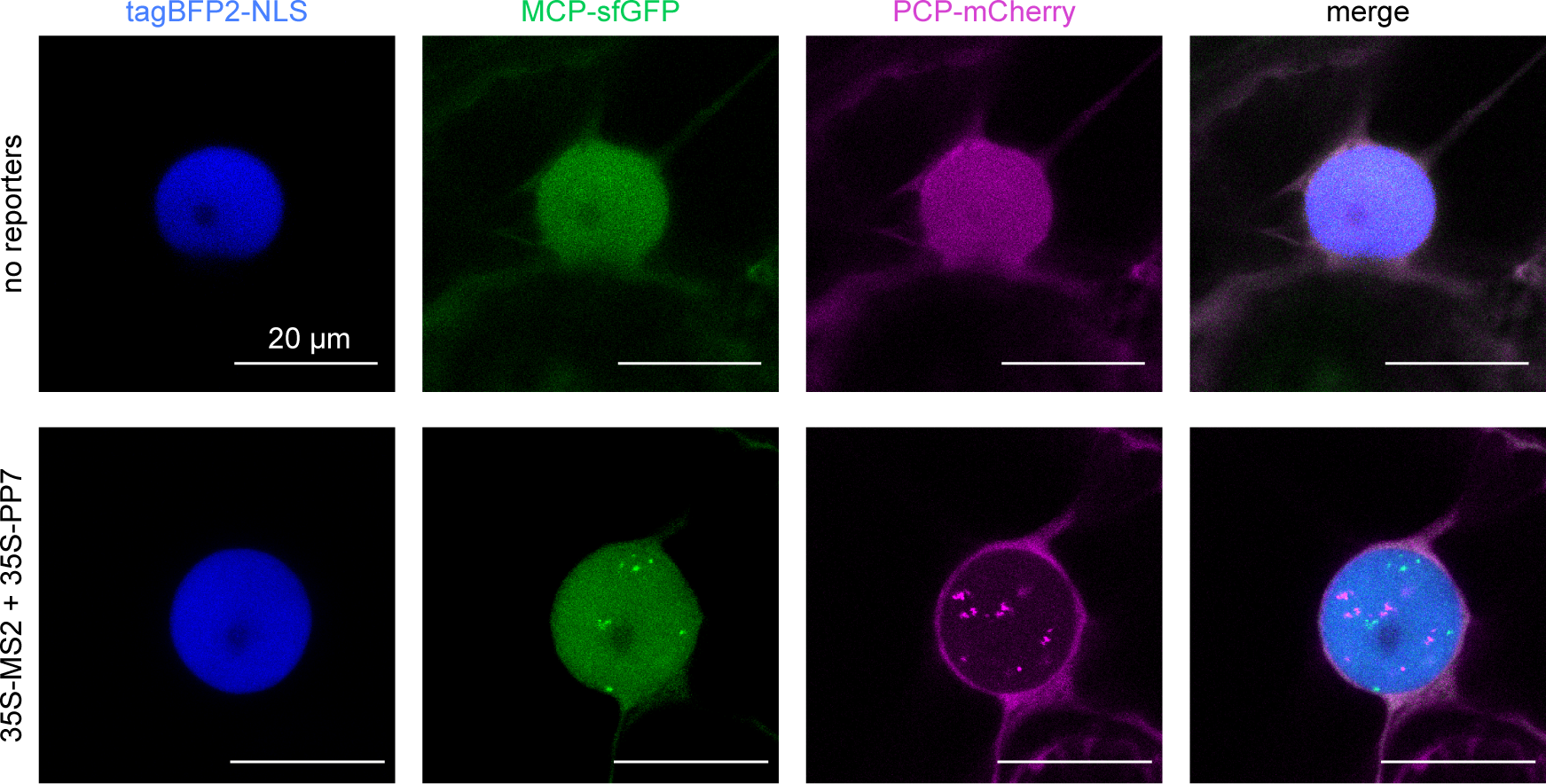
Related to Figure 1F. MCP-sfGFP and PCP-mCherry are homogeneously distributed in the nucleus in the absence of transcription. Maximum fluorescence projection snapshot of the nucleus of a Tobacco cell expressing MCP-sfGFP, PCP-mCherry and nuclear localized tagBFP2. No nuclear puncta appear in the absence of PP7 and MS2 reporters.

**Figure S2.**
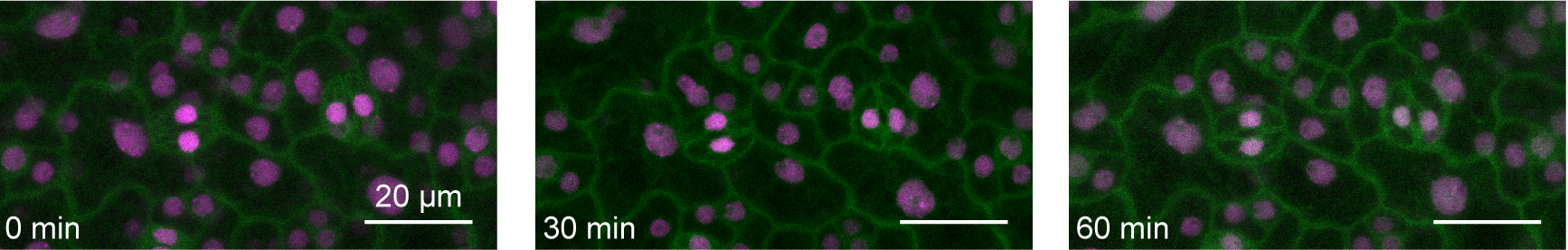
Related to Figure 2A. Lack of HSP101 induction at room temperature. Maximum z-projected image snapshots of the PCP-GFP/HSP101-PP7 Arabidopsis line imaged at room temperature. No spots were detected after continuous imaging for 80 minutes. Scale bar = 20 *µ*m.

**Figure S3.**
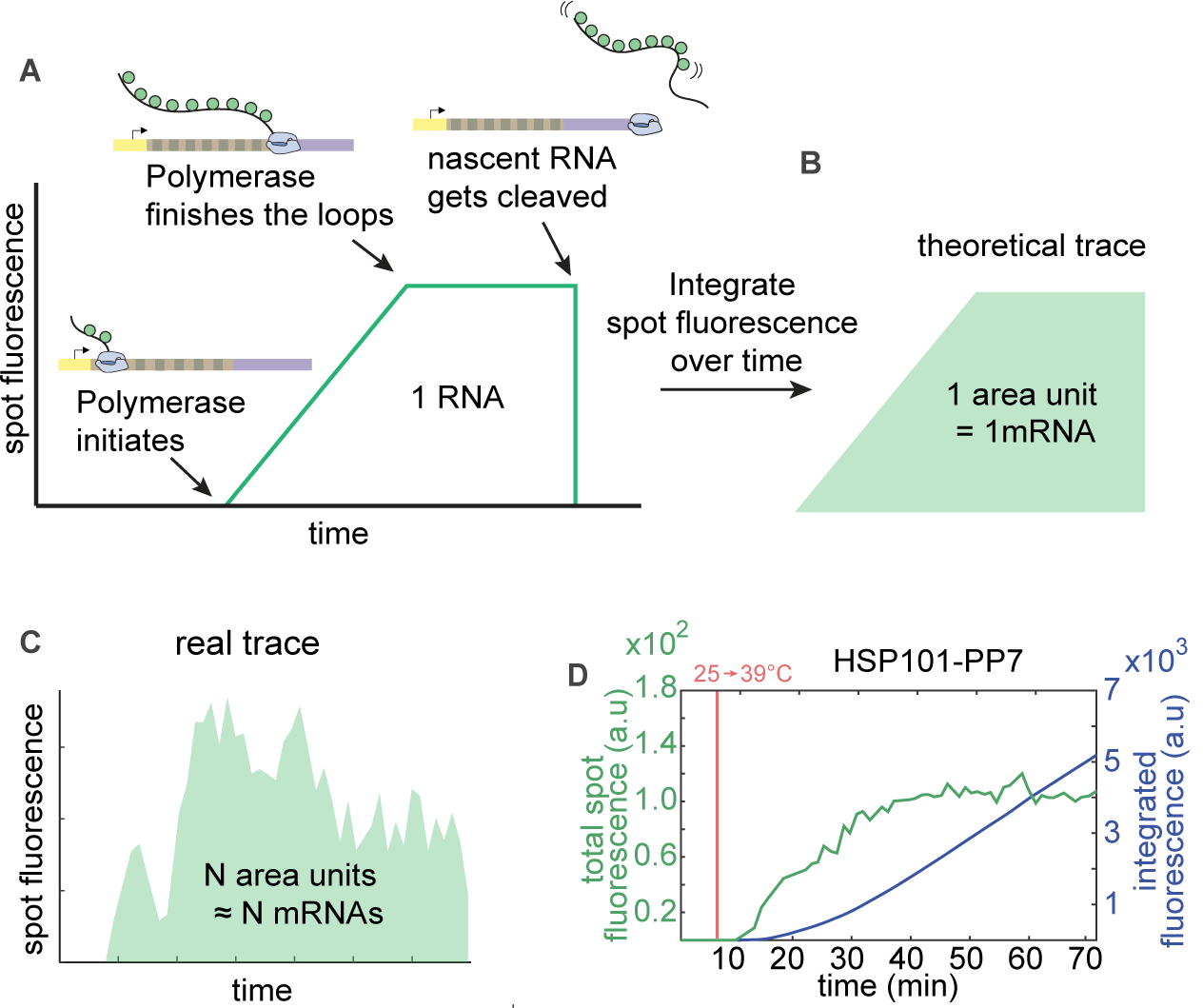
Related to Figure 2B. Integrated fluorescence as a metric for total mRNA produced. **(A)** Fluorescence profile of a single RNAP molecule as it traverses the gene. (B) Integrating this curve over time yields a unit of area associated with the production of a single mRNA molecule. (C) In the case of an actual transcription spot—resulting from the activity of multiple polymerase molecules—the integrated fluorescence over time will correspond to a number of area units equal to the number of produced mRNA molecules. **(D)** Data from a HSP101-PP7 replicate from Figure 2. Total spot fluorescence normalized by the number of cells in the field of view (green) and time integral of this signal (blue). The red horizontal line indicates when the stage temperature was shifted from room temperature to 39 °C.

**Figure S4.**
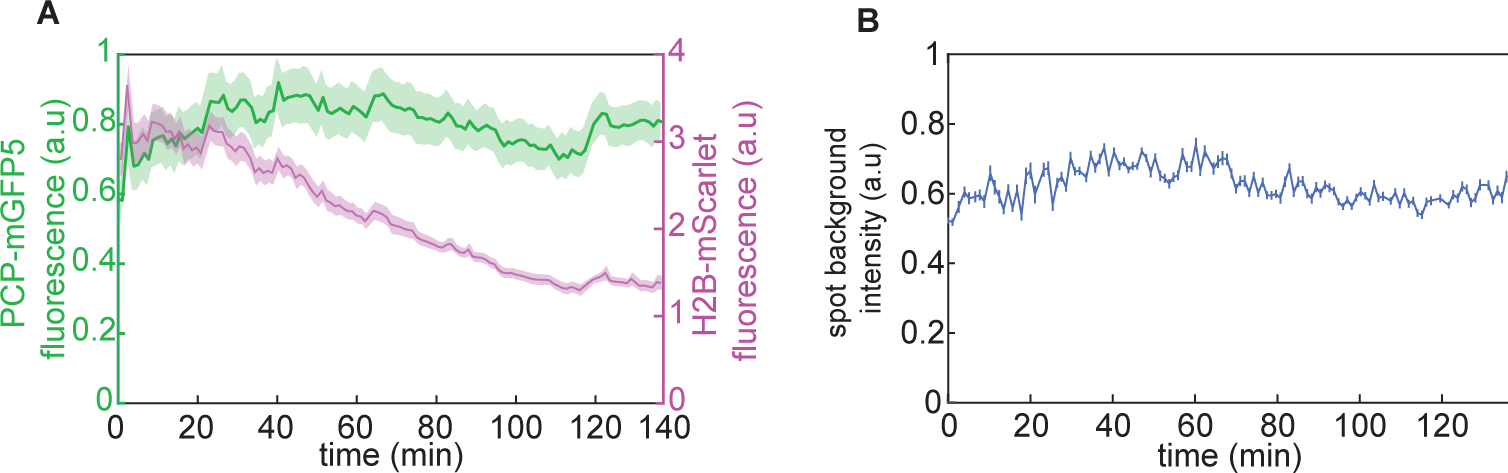
Related to Figure 2B. Control for photobleaching. **(A)** Mean and standard error of nuclear fluorescence from PCP-GFP and Histone 2B-mScarlet in an Arabidopsis line expressing PCP-GFP and a reporter construct driven by the constitutive EF-Tu promoter imaged using our standard imaging conditions. There is no significant bleaching of the PCP-GFP signal even after imaging for more than two hours. However, there is evident bleaching of mScarlet. **(B)** No bleaching is observed in the fitted PCP-GFP spot fluorescence background in the same experiment either.

**Figure S5.**
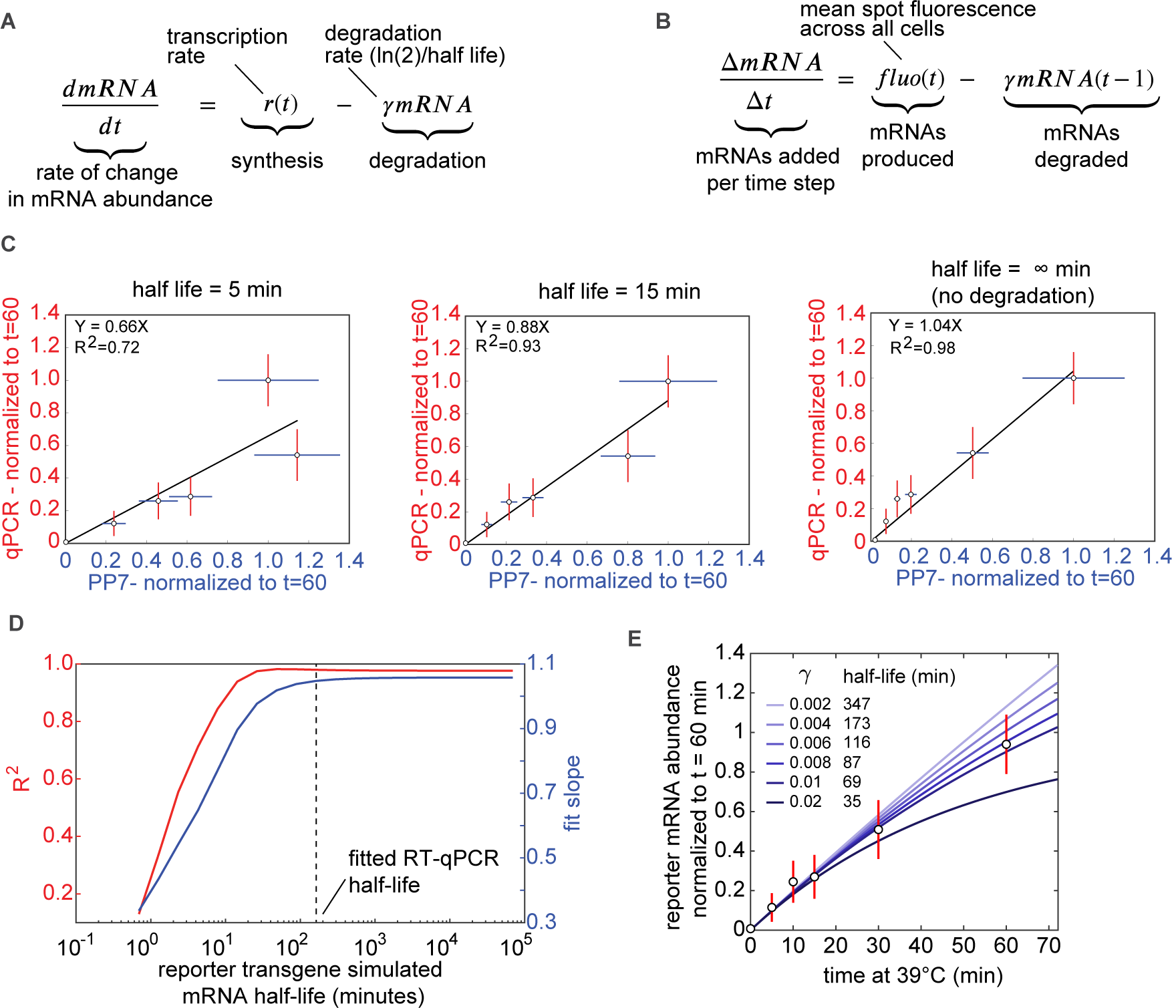
Related to Figure 2B. Exploring the effect of the mRNA degradation rate on the validation of the PP7 system against RT-qPCR measurements. **(A)** The rate of change in mRNA abundance is determined by a time-dependent rate of mRNA synthesis *r*(*t*) and a constant mRNA degradation rate *γ*. **(B)** Discretized version of equation (A) used to obtain the accumulated mRNA based on spot fluorescence measurements. At each time point the rate of synthesis is equal to the spot fluorescence while the number of mRNA molecules accumulated up to the previous time point are degraded at a simulated rate *γ*. Note that the mRNA half-life is defined as *τ*_1/2_ = ln(2)/*γ*. **(C)** Linear regression between the reporter mRNA abundance measured by RT-qPCR versus microscopy as in Figure 2C using the equation in (B) to incorporate mRNA degradation into the microscopy-based measurement. Because microscopy only reports on the synthesized, and not the degraded mRNA, we considered different, constant degradation rates and included this correction in the linear regression. **(D)** Fit parameters (*R*^2^ and fit slope) as shown in (C) were calculated for a range of mRNA degradation rates expressed as half-lives. There is a good correlation and a constant slope between RT-qPCR and microscopy for half-lives longer that ∼10 minutes. The dashed horizontal line indicates the fitted reporter mRNA half-life obtained in (C). **(E)** the reporter mRNA abundance measured by RT-qPCR was fitted to the mRNA accumulation model in (A) assuming a constant synthesis rate. mRNA accumulation according to RT-qPCR is almost linear on the timescales tested, resulting in a relatively long half-life. This half-life value is within the regime where there is a good correlation between PP7 fluorescence and qPCR (see vertical dashed line in (D)).

**Figure S6.**
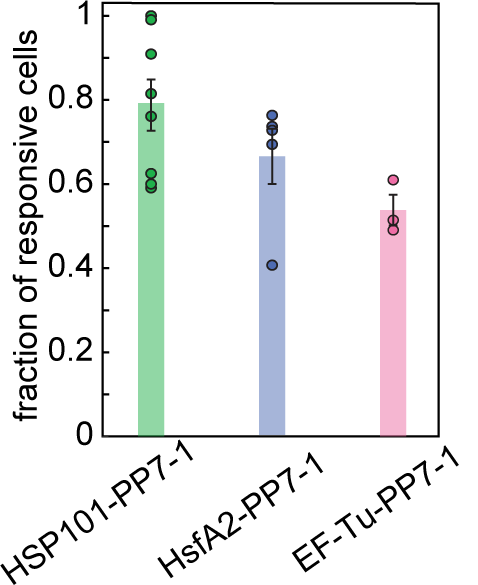
Related to Figure 3A. Reproducibility of the fraction of responsive cells. Mean and standard error of the fraction of transcriptionally responsive cells, defined as the number of nuclei that display reporter activity at least at one time point during the experiment divided by the total number of nuclei in the field of view (see Fig. 3A, bars on the right of each heat map). Circles represent at least three biological replicates

**Figure S7.**
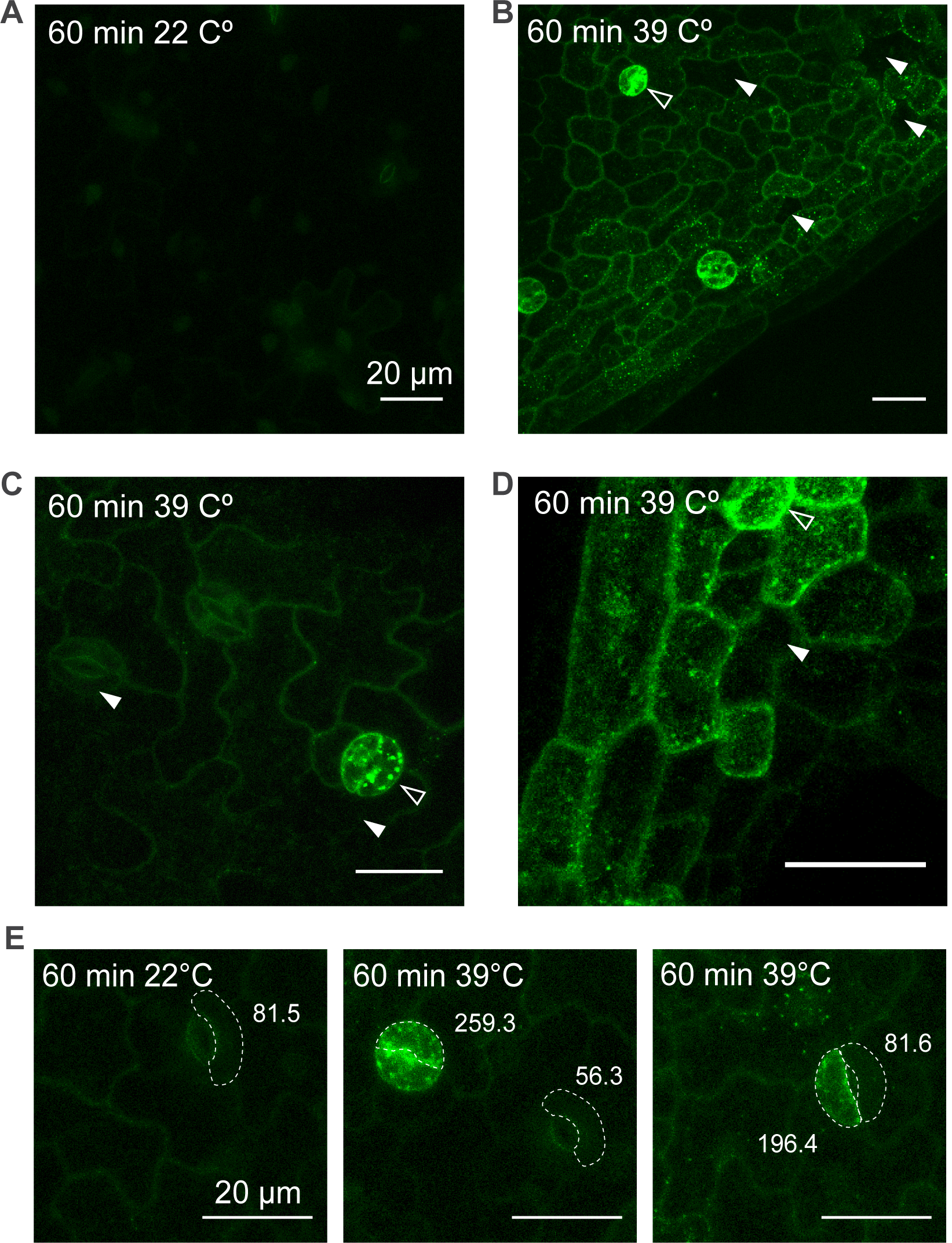
Related to Figure 3: A rescue construct of HSP101-GFP reveals how refractory cells lead to substantial cell-to-cell heterogeneity in HSP101-GFP accumulation upon heat shock. **(A-E)** Maximum fluorescence projections of leaf epidermis cells from *hsp101* knockout mutant plants complemented with a transgene coding for a HSP101-GFP fusion driven by 734 bp of the endogenous HSP101 promoter (***McLoughlin et al., 2016***). Detached leaves were treated with 39 or 22 °C for 60 minutes prior to imaging. **(A)** Untreated control. **(B-D)** Treated samples. White filled arrowheads indicate cells with negligible levels of GFP accumulation. Empty white arrowheads indicate cells with high levels of GFP accumulation. **(E)** Quantification of GFP fluorescence in treated and untreated cells. The dashed line highlights cells whose fluorescence was calculated. The numbers next to each cell correspond to the integrated GFP fluorescence of the volume of each cell highlighted.

**Figure S8.**
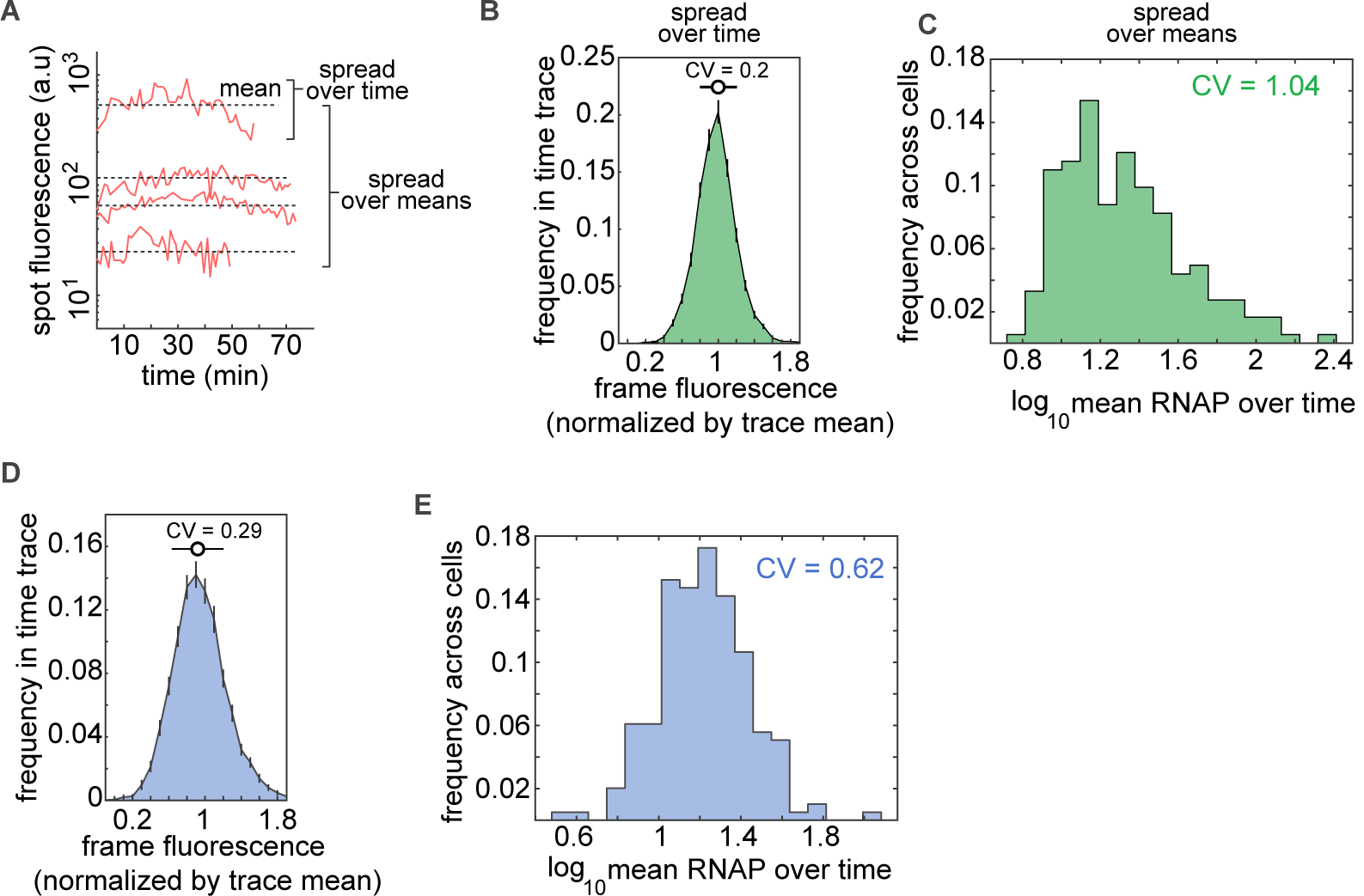
Related to Figure 4: Spot fluorescence varies widely across cells but is relatively stable over time in individual cells. **(A)** Representative spot fluorescence time traces in HSP101-PP7-1 replicates from Figure 3. Dashed lines correspond to the mean level of fluorescence of each trace over time. The spread of fluorescence values around this mean for each individual trace (“spread over time”) informs about temporal fluctuations in transcriptional activity for each individual spot (B). The variability of mean fluorescence values across cells, is captured by the “spread over means”, and informs about cell-to-cell heterogeneity in activity (C). **(B)** Distribution of frame fluorescence values normalized by the mean over time for each fluorescence trace pooled from all HSP101-PP7-1 replicates from Figure 3. The spread over time of fluorescence values of a given spot is very close to the mean, resulting in a coefficient of variation (CV=standard deviation/mean) of 0.2. **(C)** Distribution of mean fluorescence over time (see dashed lines in (A)) of all cells in HSP101-PP7-1 replicates. The average transcriptional activity varies widely across cells, with a coefficient of variation of 1.04. **(D**,**E)** Same as (B) and (C) for HsfA2-PP7-1 fluorescence traces pooled across replicates from Figure 4.

**Figure S9.**
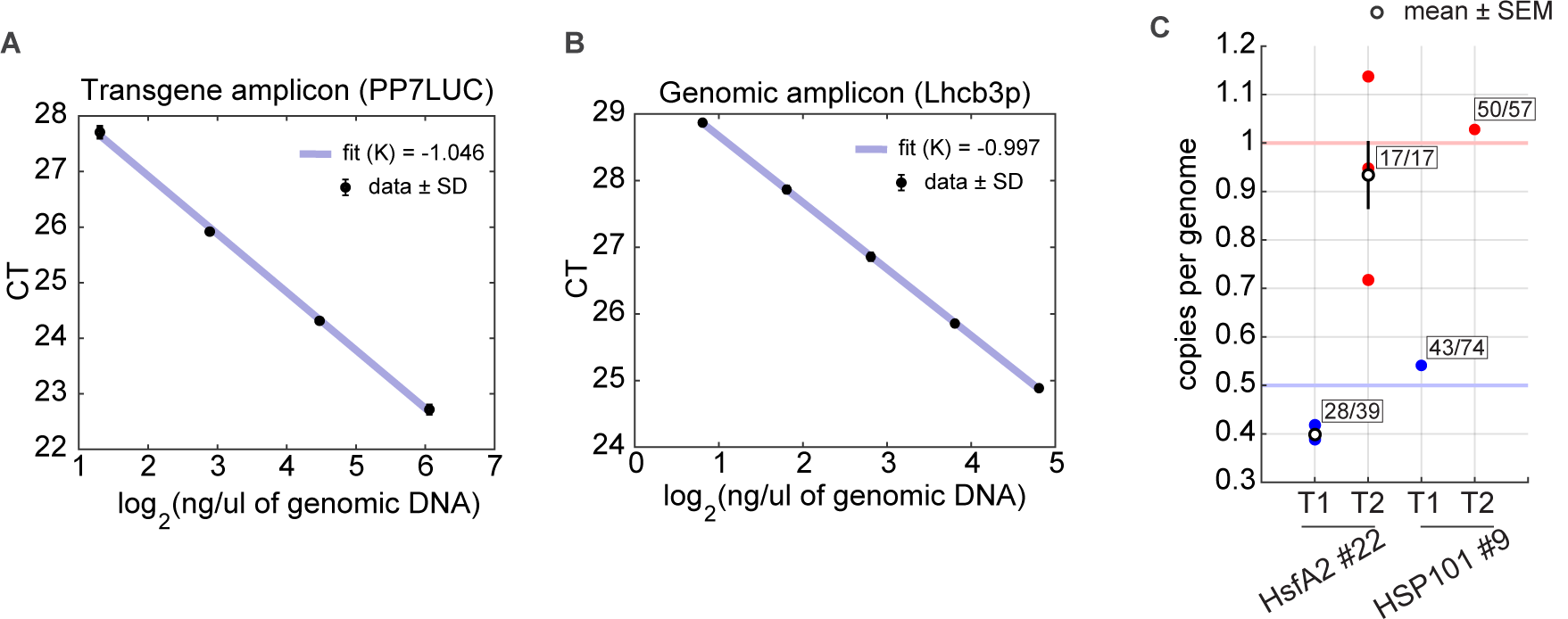
Amplification efficiency of primer pairs and determination of the copy number of single insertion lines. **(A)** qPCR results for serial dilutions of transgenic Arabidopsis plants using primer pairs targeting the reporter transgene. **(B)** Same as (A) for a primer pair targeting a genomic location upstream of the *Lhcb3* gene. In (A) and (B), the slope of the linear fit corresponds to *K* = 1/(1 + *log*_2_(*ϵ*) where *ϵ* is the amplification efficiency. **(C)** Number of copies of the PP7 reporter transgene per genome copy in two single insertion reporter lines in the T1 and T2 generations. The horizontal blue line indicates the expected value for a single-copy hemizygous plant where the insertion locus contains a single copy of the transgene. The red horizontal line indicates the expected value for a plant homozygous for a single insertion where this insertion contains a single copy of the transgene. The ratios next to each data point indicate the fraction of survivors over the total number of plated seeds under kanamycin selection.

**Figure S10.**
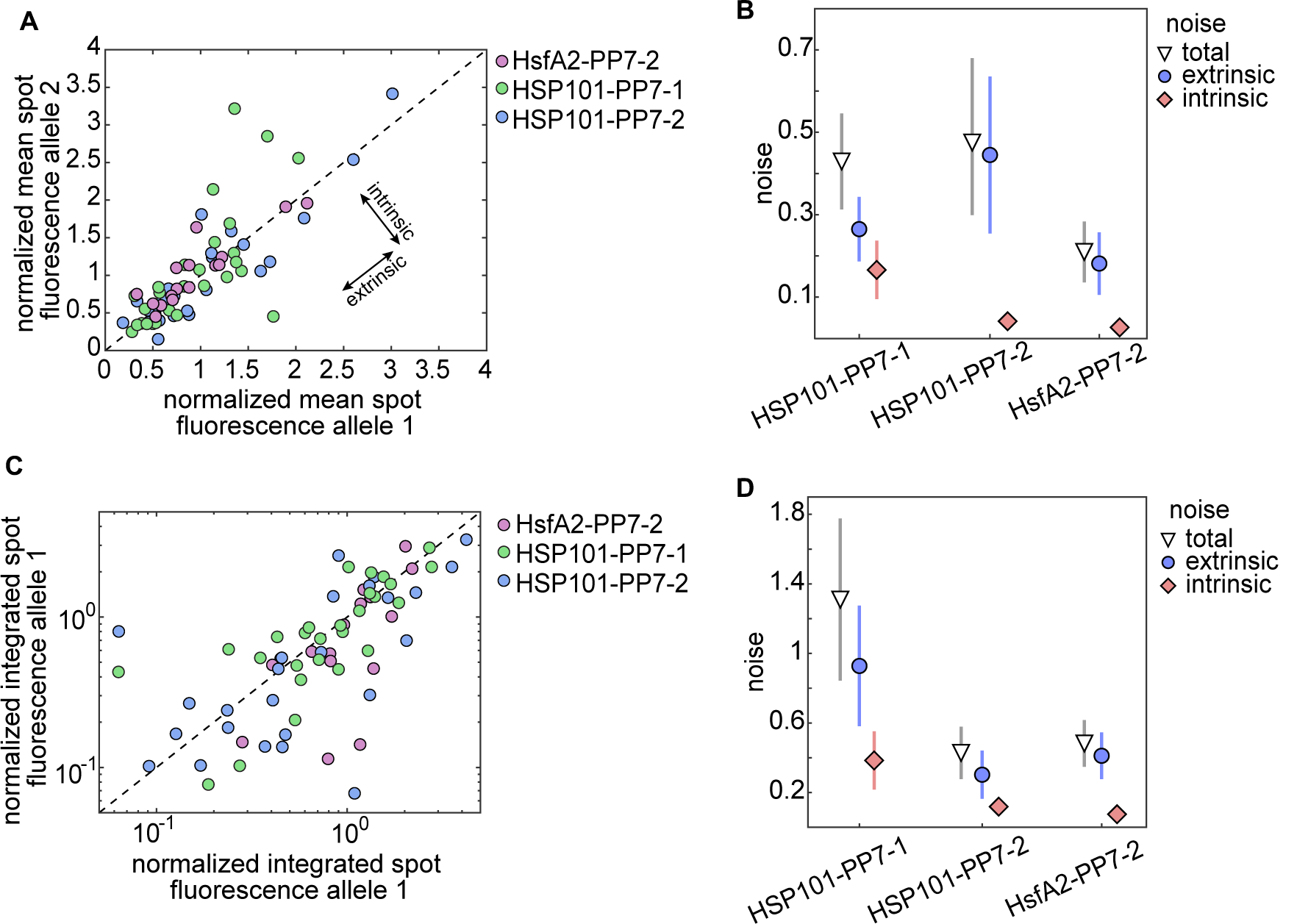
Related to Figure 5 Extrinsic noise is larger than intrinsic noise among nuclei with two active alleles. **(A)** Scatter plot showing the mean spot fluorescence over time for allele pairs belonging to the same nucleus in three different single-insertion lines homozygous for the PP7 reporter. **(B)** Decomposition of the total variability in (A) into its intrinsic and extrinsic components. **(C)** Scatter plot of integrated fluorescence over time in allele pairs belonging to the same nucleus in three different single-insertion reporter lines homozygous for the PP7 transgene (same as Figure 5E except that inactive alleles are not included). **(C)** Decomposition of the total noise in (C). In (A) and (C) values were normalized to the mean across all alleles in that line and the diagonal line shows y=x. Error bars in (B) and (C) correspond to the bootstrapped error.

### S4 Supplementary Videos

S1. **Video 1. Constitutive reporter in tobacco**. Movie of tobacco cell expressing PCP-GFP and GAPC2-PP7. The scale bar is 10 *µ*m.

**Video 2. Inducible reporter in tobacco**. Movie of tobacco cell expressing PCP-GFP and HSP70-PP7 under heat shock treatment starting at 10 min. The scale bar is 10 *µ*m.

**Video 3. Inducible HSP101-PP7 reporter in Arabidopsis tissue**. Movie of leaf cells in Arabidopsis line stably transformed with PCP-GFP and HSP101-PP7 under heat shock treatment starting at 6 min. The scale bar is 10 *µ*m.

**Video 4. Inducible HsfA2-PP7 reporter in Arabidopsis tissue**. Movie of leaf cells in Arabidopsis line stably transformed with PCP-GFP and HsfA2-PP7 under heat shock treatment starting at 8 min. The scale bar is 10 *µ*m.

**Video 5. Constitutive reporter in Arabidopsis tissue**. Movie of leaf cells in Arabidopsis line stably transformed with PCP-GFP and EF-Tu-PP7. The scale bar is 10 *µ*m.

**Video 6. Arabidopsis plant homozygous for an inducible reporter**. Movie of leaf cells in a homozygous Arabidopsis line stably transformed with PCP-GFP and HSP101-PP7 under a heat shock treatment starting at 0 min. The scale bar is 10 *µ*m.

